# Impact of selective logging on the population dynamics and genetic diversity in the neotropical timber tree *Dicorynia guianensis*

**DOI:** 10.1101/2025.09.26.678763

**Authors:** Julien Bonnier, Stéphane Traissac, Olivier Brunaux, Valérie Troispoux, Niklas Tysklind, Myriam Heuertz

## Abstract

Selective logging is a widely applied forest management strategy in the tropics, yet it is insufficiently documented how it affects the genetic and demographic processes of seedling recruitment in timber species in the Guiana Shield. Our study investigates how selective logging influences genetic diversity, gene dispersal, and seedling establishment in *Dicorynia guianensis* by comparing a logged and an unlogged forest plot in French Guiana. We genotyped 703 individuals using 66 nuclear SSR markers and applied parentage analyses to infer dispersal patterns and reproductive success. We analysed genetic diversity and spatial genetic structure across life stages, and tested whether seedling recruitment was associated with logging-related canopy openings. Genetic diversity indices were broadly similar between logged and unlogged plots, with no evidence of genetic erosion in adults or seedlings. Seedling establishment was associated with logging-induced canopy openings. Parentage analyses revealed shorter mean dispersal distances in the logged plot but substantial long-distance pollen flow, ensuring admixture and inter-plot connectivity. Reproductive success was more evenly distributed among mothers, whereas male contributions were skewed in the logged plot. Selective logging did not cause immediate genetic erosion but altered dispersal dynamics and reproductive patterns. These findings underline the resilience of *D. guianensis* under current management practices, while emphasizing the need for long-term monitoring of regeneration to ensure sustainable recruitment and evolutionary resilience across logging cycles.

## 1. Introduction

Tropical forests, and particularly the Amazon rainforest, are increasingly subject to selective logging as part of reduced-impact forest management strategies aimed at reconciling timber production with forest conservation. Unlike clear-cutting, selective logging removes only a fraction of canopy trees, typically the largest individuals of commercial species, while attempting to maintain forest structure and ecological processes. Evidence suggests that even low-intensity logging can lead to significant alterations in forest composition, biomass, and ecosystem functioning (Asner et al., 2009). In the Amazon, these operations have expanded rapidly, while the cumulative effects of logging, road construction, and increased hunting pressure remain poorly quantified (Putz et al., 2000, 2008). While selective logging is promoted as a more sustainable alternative to deforestation, the long-term impacts on ecological and evolutionary processes, including tree regeneration, genetic diversity, and reproductive dynamics are still subject to uncertainty. This is particularly true in the Guiana Shield, where timber exploitation is recent and highly selective, yet the resilience of tree populations to disturbance remains largely unexplored.

In French Guiana, selective logging is the primary mode of timber extraction, governed by the “Office national des Forêts” (ONF) regulations that aim to ensure sustainable management of French Guiana tropical rainforests. The current system advocates the removal of a limited number of commercial trees per hectare (4 - 5 stems per hectare, every 65 years, minimum cutting diameter: 55 cm DBH) (Guitet et al., 2014; PRFB, 2019). Despite this low extraction intensity, single logging cycles can substantially modify forest structure, reduce the abundance of large reproductive individuals, and alter local microhabitats. Logging gaps and skid trails not only create heterogeneity in light and soil conditions but also increase accessibility for hunting, thereby indirectly affecting animal-mediated seed dispersal. While this system is often promoted as a model of low-impact logging, particularly when compared to practices in other parts of Amazonia, the long-term consequences for tree population viability remain insufficiently assessed. In particular, little is known about how such interventions affect the regeneration dynamics, reproductive success, and genetic composition of exploited tree species over time (Wernsdörfer et al., 2011).

Selective logging, even at low intensity, can have implications for the evolutionary dynamics of tropical tree populations by altering both demographic and genetic processes. While adult genetic diversity may appear stable in the short term, disturbances such as tree felling and canopy openings can shift mating patterns, increase spatial isolation among reproductive individuals, and reduce effective population size (de Oliveira et al., 2020; Leclerc et al., 2015). Unlike temperate forests, where tree species densities are typically higher, tropical tree species often occur at low densities, making their reproductive systems particularly sensitive to reductions in adult abundance (Finkeldey & Ziehe, 2004). Empirical studies have reported reductions in allelic richness and elevated selfing rates in logged populations, particularly when large trees, often the primary pollen donors, are selectively removed (Duminil, Mendene Abessolo, et al., 2016). The preferential removal of fast-growing or well-formed individuals can induce dysgenic selection, compromising the genetic quality of future generations (Jennings et al., 2001). Post-logging regeneration represents a critical stage during which genetic and demographic shifts become most apparent. It can result in regenerated cohorts that are genetically less diverse and more spatially aggregated than those in undisturbed conditions, with shallower spatial genetic structure and increased relatedness among individuals (de Oliveira et al., 2020; Leclerc et al., 2015; Wernsdörfer et al., 2011). These effects are compounded when logging is coupled with changes in microhabitats, pollinator activity, or seed dispersal agents, leading to assortative mating and potential inbreeding depression (Jennings et al., 2001; Loveless & Hamrick, 1984; Monthe et al., 2017). The resulting erosion of within-population genetic diversity ultimately compromises the evolutionary potential, adaptability, and long-term resilience of exploited tree species (Fargeon et al., 2016).

Tree regeneration is often concentrated in felling gaps and skid trails, where light and soil disturbance create favourable microsites for seedling establishment (Fredericksen & Mostacedo, 2000; Swaine & Agyeman, 2008). In theory, canopy openings created by felling operations can enhance light availability and reduce competition, thereby facilitating recruitment and growth of residual or new individuals (Swaine & Agyeman, 2008). However, this regeneration may be localized and genetically clustered, especially in species with limited seed dispersal, resulting in overrepresentation of the nearest fruiting trees and increased spatial genetic structure in post-logging cohorts (Hardy et al., 2019; F. A. Jones et al., 2005; Sezen et al., 2005). Gene flow, through seed and pollen dispersal, plays a key role in mitigating these risks by introducing genetic novelty, maintaining heterozygosity, and enabling adaptation to environmental changes (Levin & Kerster, 1974; Monteiro et al., 2019; Wernsdörfer et al., 2011). Selective logging can significantly disrupt mating patterns and reduce the effective gene flow within and between populations (Duminil, Mendene Abessolo, et al., 2016; Silva et al., 2008). Understanding how these processes interact under logging regimes is thus essential for predicting the evolutionary trajectory and regeneration success of exploited tree species.

*Dicorynia guianensis* Amshoff (Fabaceae), locally known as Angélique, is an ecologically and economically important endemic tree species of the Guiana Shield. It is among the 16 hyperdominant species of the Amazon basin in terms of wood biomass and productivity (Fauset et al., 2015), and represents over 50% of the timber production in French Guiana (PRFB, 2019). The species occurs predominantly in terra firme lowland rainforests and thrives on clayey or sandy soils, with a preference for well-drained sites (Falcão et al., 2022; Guitet et al., 2014). *D. guianensis* occurs in aggregated spatial distributions, typically forming genetically differentiated clusters ranging from four to 18 reproductive individuals per hectare (Bariteau, 1993; Kokou, 1994; Latouche-Hallé et al., 2003). The species is pollinated by large bees and is predominantly outcrossing, with average pollen dispersal distances of 200 meters (Caron et al., 1998; Forget, 1988). Seed dispersal is mostly gravity- and wind-mediated, resulting in short dispersal distances rarely exceeding 50 meters (Jésel, 2005). Seeds are subject to predation by insects, parrots, and rodents, with limited evidence of effective secondary dispersal (Bonnier et al., 2026; Jésel, 2005). The combination of limited seed dispersal, low population densities, and selective logging targeting the largest reproductive trees renders *D. guianensis* particularly vulnerable to demographic and genetic erosion. Previous genetic studies have revealed substantial spatial genetic structure and east-west differentiation in French Guiana, based on both nuclear and chloroplast markers (Bonnier et al., 2023, 2025; Caron et al., 2000). As such, *D. guianensis* provides an ideal model to investigate impacts of logging on regeneration processes, reproductive success, and gene flow in tropical forest trees.

Here, we assess the genetic and demographic consequences of selective logging in *D. guianensis* by comparing two nearby natural forest plots in French Guiana: one unlogged (HKO50) and one selectively logged in 2012 (PAI74). By focusing on an ecologically and economically important species, our aim is to provide an integrative perspective on how selective logging in French Guiana can affect forest resilience. We combine spatially explicit genetic data and demographic metrics to address the following questions: 1) Does selective logging affect the genetic diversity and spatial genetic structure of *D. guianensis* populations? 2) To what extent does logging influence gene flow dynamics and reproductive success? 3) What are the consequences of canopy gaps on seedling recruitment and genetic composition? By addressing these questions, we aim to provide empirical insights into how forest management practices influence the evolutionary trajectory and regeneration capacity of a key timber species in the Guiana Shield. Our findings contribute to the broader understanding of sustainable tropical forest management and emphasize the value of integrating demographic and genetic metrics into long-term monitoring frameworks.

## 2. Materials and method

### 2.1. Sampling and forest plots

Our study was conducted in two forest plots located in the Régina-Saint Georges forest massif in eastern French Guiana (Figure 1). The area is currently being logged, which allows better access to the forest through the creation of tracks, increasing hunting pressure. The plots are embedded within lowland terra firme rainforests dominated by *D. guianensis*, and are part of the DYGEPOP research network established to monitor the population dynamics and silvicultural responses of high-value timber species in managed and unmanaged tropical forests. The first plot, PAI74, covers 17 hectares and was selectively logged in 2012 as part of a silvicultural trial. Forty-two reproductive trees with a DBH ≥50 cm were harvested across the plot, following standard forestry guidelines. Notably, a first sampling of adult trees was conducted in 2009 for the DYGEPOP project, prior to logging operations, enabling a rare opportunity to assess the genetic composition of reproductive individuals before tree extraction. Spatial mapping of disturbance features such as skid trails and canopy gaps have been mapped after logging. A second field campaign in 2023 allowed for the sampling of all remaining adult trees, as well as juvenile and intermediate individuals (≥1 cm DBH). In total, 418 individuals of *D. guianensis* were sampled in the PAI74 plot.

**Figure 1.**
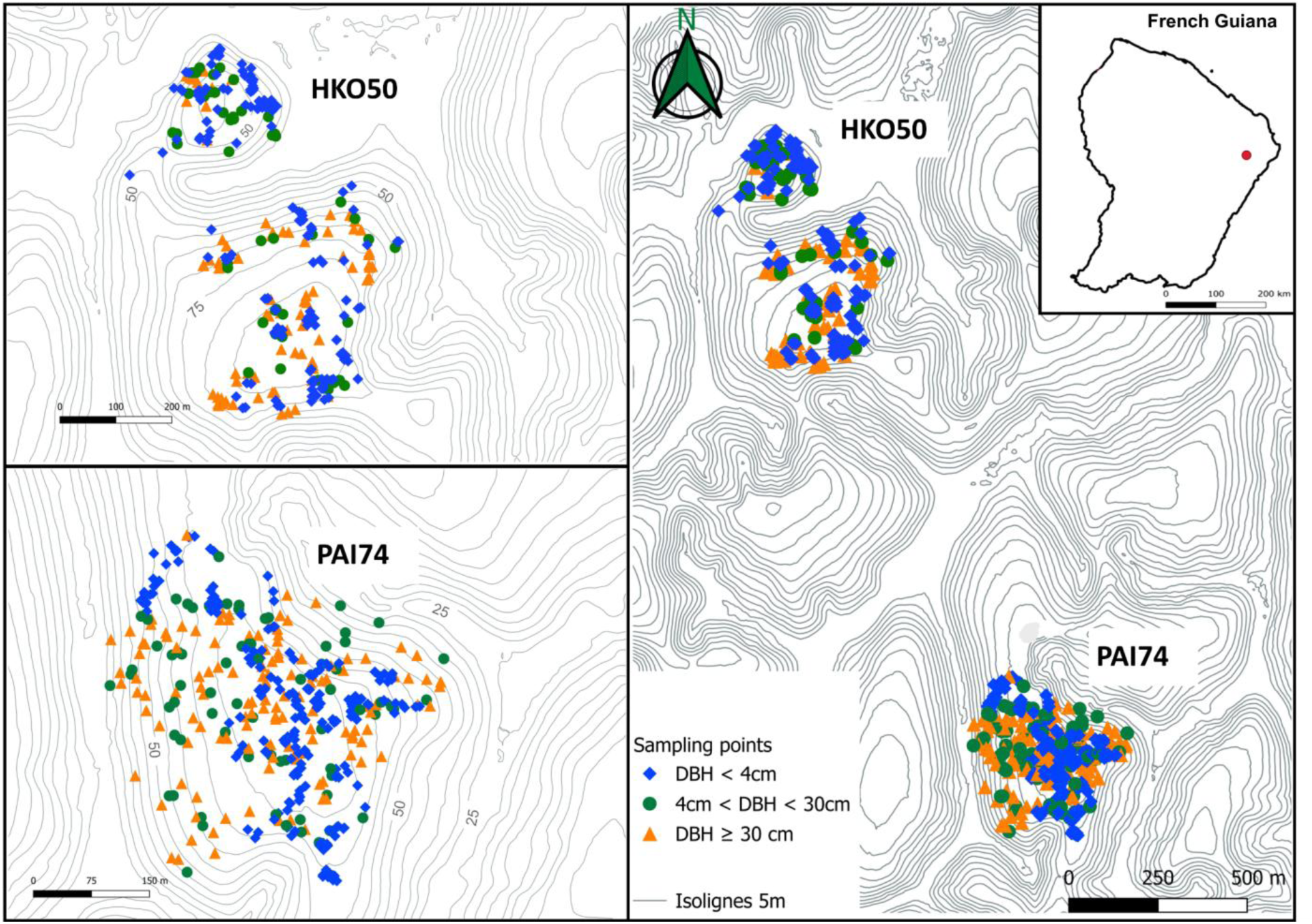
Geographic location and spatial distribution of sampled *Dicorynia guianensis* individuals across two study plots in French Guiana. Map showing the spatial distribution of *D. guianensis* individuals in the unlogged plot HKO50 and the selectively logged plot PAI74. Points represent individual trees, categorized by diameter at breast height (DBH): seedlings (DBH < 4 cm, blue diamonds), saplings (4 cm ≤ DBH < 30 cm, green circles), and adults (DBH ≥ 30 cm, orange triangles). Topographic isolines (5 m elevation intervals) illustrate the relief across both plots.

The second plot, HKO50, spans 21 hectares and serves as an unlogged control, representing undisturbed population dynamics in the same forest massif (1.5 km away from PAI74). No silvicultural intervention has occurred within HKO50, and the site provides a natural benchmark against which to evaluate the impacts of logging. A complete inventory of *D. guianensis* individuals ≥1 cm DBH was carried out in 2024, resulting in the sampling of 285 individuals distributed across life stages.

In both plots, all sampled individuals were georeferenced using GPS, the elevation of their position and their diameter at breast height (DBH) were recorded. Leaf or cambium samples were collected and preserved in silica gel under controlled room temperature conditions until DNA extraction. Due to high growth rate variability in tropical trees, precise age classes cannot be reliably defined. Instead, individuals were categorized into three DBH-based life-stage cohorts reflecting ecological and reproductive status: juveniles (JUV: <4 cm DBH), representing the most recent generations established after the logging operations conducted 12 years prior to our sampling campaign in PAI74; adults (ADL: >30 cm DBH), representing reproductively mature individuals capable of contributing to future generations as determined by field observations of flowering and fruiting behavior; and intermediates (INT: 4-30 cm DBH), representing individuals that are neither recent recruits nor reproductively active. For adults (>30 cm DBH), complete sampling was achieved through comprehensive forest inventories conducted by the Office National des Forêts (ONF). For smaller individuals (<30 cm DBH), we aimed for exhaustive sampling with intensive ground searches, but acknowledge that some individuals may have been missed due to detection limitations in dense understory conditions. We estimate that fewer than 10% of individuals in the smaller size classes (<30 cm DBH) were not detected during our surveys, considering that the samples were collected by specialists of the species over the span of a week of sampling.

### 2.2. DNA isolation, genotyping, and contamination

Genomic DNA was extracted from 50 mg of silica-dried leaf or cambium tissue using a CTAB-based protocol following (Doyle & Doyle, 1990). Each sample was individually ground into a fine powder using a TissueLyser (Qiagen), and DNA quantity and quality were assessed using a NanoDrop 8000 spectrophotometer (Thermo Fisher Scientific, MA, USA). Extractions yielding DNA concentrations below 30 ng/μL were repeated until an acceptable quality was obtained.

Microsatellite genotyping was conducted using two nuclear and one chloroplast SSR marker panels previously developed for *D. guianensis* and described in Bonnier et al. (2026). The markers were genotyped using a high-throughput SSR sequencing approach (SSR-SEQ) on Illumina sequencers; this multiplexed sequencing method enabled cost-effective processing of all loci across individuals while maximising parentage assignment power and reliability in this spatially complex tropical forest system. While the marker panels and genotyping protocol remained identical to that used in Bonnier et al. (2026), a different encoding scheme was applied for allele representation. A total of 66 nuclear SSR markers and 26 chloroplast SSR markers were retained for subsequent analyses. Samples with over 25% missing nuclear data were removed from the analysis (n=5); this threshold was applied as a uniform quality filter across all analyses (genetic diversity, SGS, and parentage assignment) to ensure comparability and minimize biases introduced by unequal genotyping success.

To assess potential contamination or sample duplication, multilocus genotypes and multilocus lineage were identified using the MLG_tab and MLL_generator functions of the RClone ver 1.0.3 package (Arnaud-Haond et al., 2007). Genetic distance distributions between observed multilocus genotypes were compared to those generated from simulated datasets in RClone (with and without selfing), in order to provide a reference baseline and identify unusually low divergence values indicative of potential contamination or clonal replication. Multilocus Lineage (MLL) was identified using a genetic distance threshold (alpha2 = 2), allowing grouping of genetically redundant or closely related individuals. Genotypic diversity and clonal structure were evaluated using clonal_index and the Pareto index (Pareto_index). Null allele analysis was conducted using the PopGenReport package (Adamack & Gruber, 2014) implementing Brookfield (1996) estimators. Bootstrap confidence intervals (n=1000) were calculated to assess the reliability of null allele frequency estimates. All filtering, quality control, and contamination detection steps were carried out prior to downstream genetic analyses, ensuring the reliability of the genotype dataset.

### 2.3. Population structure

Bayesian clustering analysis was performed using STRUCTURE v2.3.4 (Pritchard et al., 2000) to identify the number of genetic clusters among the two plots and assess the degree of admixture of individuals. Analyses were run across a number of clusters, *K,* ranging from 1 to 10, using a burn-in period of 50,000 iterations followed by 500,000 MCMC repetitions, with ten replicates per *K*. STRUCTURE Harvester (Earl & vonHoldt, 2012) was used to determine the most likely number of clusters using the variation of LnP(*K*) as a function of *K*, and the ΔK method (Evanno et al., 2005). SPAGeDi software v1.5a (Hardy & Vekemans, 2002) was used to compute genetic differentiation with *F*_ST_ ^(^Weir & Cockerham, 1984^)^ between the two plots. Significant differentiation (*F*_ST_) was assessed with 10 000 permutation of individuals between plots.

To further assess spatial patterns of genetic variation, spatial principal component analyses (sPCA) were conducted using the *spca()* function in the R package adegenet (Jombart, 2008). sPCA was applied independently within each plot, and jointly across both plots to detect spatially structured genetic gradients. Global and local Monte Carlo tests were used to test the significance of spatial patterns. The global test (G-test) evaluates large-scale spatial genetic structure, or spatial autocorrelation throughout the plot, while the local test (L-test) detects high genetic divergence in nearby individuals. Spatial interpolation of lagged scores was used to visualize the spatial distribution of genetic variation across the study area. Principal component analysis (PCA) were analysed with adegenet to visualize patterns of genetic differentiation between the two forest plots and among cohorts within and across plots.

### 2.4. Genetic diversity and spatial genetic structure

Genetic diversity and spatial genetic structure were assessed using nuclear microsatellite data. Analyses were conducted independently on the two plots and for the three DBH-based cohorts (JUV, INT, ADL). For PAI74, genetic diversity was computed both for the current post-logging adult population (n = 93), and for the full pre-logging adult dataset (n = 135), taking advantage of archived plant material collected before logging. Diversity indices were calculated using SPAGeDi v1.5a (Hardy ^&^ Vekemans, 2002^)^. The following metrics were computed: allelic richness (*A*_R_) standardized to 30 gene copies, effective number of alleles (*NA*_e_), expected heterozygosity (*H*_e_), observed heterozygosity (*H_O_*), and inbreeding coefficient (*F*_IS_). The significance of *F*_IS_ was evaluated through 10,000 permutations of alleles among individuals within each plot or cohort. The selfing rate (S) was estimated from the multilocus standardized identity disequilibrium (Hardy, 2016), significance was assessed by permuting single-locus genotypes randomly among individuals within population independently for each locus (10,000 permutations). In addition, private allelic richness (*pA*_R_) was estimated using ADZE v1.0 (Szpiech et al., 2008), based on a standardized sample size of 30 gene copies. At the plot level, *pA*_R_ was calculated with HKO50 and PAI74 full plots to allow inter-plot comparisons. At the cohort level, *pA*_R_ was estimated within each plot across cohorts (SED, INT, ADL) to assess changes across life stages.

To quantify the intensity of spatial genetic structure (SGS), pairwise kinship coefficients (*F_ij_*) were calculated according to the method of Loiselle et al. (1995), and regressed against the logarithm of geographic distance. Analyses were done using nine distance classes (30, 60, 90, 130, 170, 220, 300, 600, and 1000 m), selected based on the spatial distribution of individuals and known dispersal distances of *D. guianensis*. The Sp statistic, defined as Sp = –b / (1 – *F*_1_) (Vekemans & Hardy, 2004), where b is the slope of the regression of *F*_ij_ on log distance and F*_1_* is the mean kinship within the first distance class, was used to quantify the strength of SGS. The significance of the regression slope underlying the Sp statistic was tested by permuting individual geographic locations among all individuals (10,000 permutations). For nuclear data, Wright’s neighborhood size (NS = 1/Sp) was also calculated as a proxy of the effective number of breeding individuals contributing locally to gene flow (Nunney, 2016; Wright, 1943). The significance of the variations observed between the diversity indices (*A*_R_, *H*_E_, *H*_O_, *F*_IS_, Sp, s) was tested between plots and cohorts by bootstrapping (n= 10,000). For PAI74, the post-logging population of ADL was considered in comparison with HKO50.

### 2.5. Parentage analysis

Parentage was inferred using the combined dataset from both plots using CERVUS v3.0.7 (Marshall et al., 2003) and COLONY (O. R. Jones & Wang, 2010) to ensure robust assignment of parent-offspring relationships. CERVUS uses a likelihood-based method, with simulations (10,000 offspring) run under the following parameters: 500 candidate parents, 25% sampling rate, and polygamy in both sexes. Only assignments with the relaxed (80%) or strict (95%) confidence thresholds based on the log-likelihood ratio were retained, and each analysis was repeated 10 times for consistency. COLONY was run in full-likelihood mode assuming diploid, monoecious individuals, and polygamy. The candidate parent pool was also set to 25%. Results included sibship family clusters (BestCluster file) for further family structure analysis. In COLONY, sibship clusters are obtained by evaluating all possible family configurations and selecting the one with the highest likelihood, grouping individuals into full-sib or half-sib families that best explain the observed multilocus genotypes. Parentage results from both programs were compared to identify consistent assignments. When a parent pair was found compatible with an offspring, the parent sharing the plastid haplotype with the offspring was assigned as the mother, and the other as the father. Only unambiguous haplotype matches were retained. Distances between offspring and assigned parents were calculated, distinguishing seed and pollen dispersal.

### 2.6. Dispersal dynamics

Parentage assignment results were used to quantify reproductive success and dispersal in the two study plots. Links between offspring and their assigned parents were mapped to estimate seed and pollen dispersal distances and to assess inter-plot connectivity. Spatial visualizations were produced using ggplot2 (Wickham, 2009), sf (Pebesma, 2018) and ggspatial (Dunnington et al., 2025) packages in R. Data manipulation and integration were performed with dplyr (Wickham et al., 2023), and tidyr (Wickham et al., 2014) packages in R.

For each mother and father in both plots, we quantified the number of assigned offspring. Seed dispersal distances (maternal) and pollen dispersal distances (paternal) were then compared between the two plots using Wilcoxon rank-sum tests (Wilcoxon, 1945). Inequality in reproductive output was assessed with the Gini index (Gastwirth, 1972), calculated on the distribution of offspring per parent with the ineq R package (Zeileis & Kleiber, 2014). Gini values were computed independently for mothers and fathers in each plot. The Gini index summarises the degree of inequality, ranging from 0 (all parents contribute equally) to 1 (all offspring come from a single parent), providing a synthetic measure of reproductive skew. Bootstrap resampling was implemented to test the significance of differences in Gini values between plots. In addition, the effective numbers of mothers (Ne_m) and fathers (Ne_f) were estimated from the parentage assignment results (Crow & Kimura, 1970). The effective number of parents was computed as Ne = 1 / Σpi², where pi = ni / N is the proportion of offspring assigned to parent i among all N successfully assigned offspring (Husband & Barrett, 1992). The influence of tree size on reproductive dynamics was tested separately for mothers and fathers in each plot using linear models (*lm* function, R base package). Model significance was assessed with F-tests. All was analysed in R version 4.4.3 (R Core Team, 2023).

To characterize the shape and scale of seed and pollen dispersal kernels, we used NMπ v2 software (Chybicki, 2018; Chybicki & Burczyk, 2010). Analyses were run separately for each plot with 200,000 MCMC iterations, both seed and pollen dispersal were modelled using exponential-power kernels, chloroplast haplotype data were used to distinguish seed from pollen dispersal events. Kernel shape parameters (bs and bp) were fixed at 1 (exponential kernel) as preliminary runs indicated convergence issues when freely estimated.

### 2.7. Regeneration in canopy openings after logging

To assess whether logging-induced canopy openings influenced the spatial pattern of seedling recruitment, we used the sf package (Pebesma, 2018) to analyse spatial intersections and to standardize spatial vector data in R. Three types of logging canopy features mapped during the DYGEPOP project were considered: felling gaps, skid trails, and log landings. Non-random seedling establishment in disturbed areas was assessed with a two-sample proportion test (prop.test()) followed by a fitted Firth logistic regression model (Firth, 1993) using the *logistf* package (Heinze et al., 2003). The model tested the combined effects of the three canopy features on seedling occurrence. We also examined the proximity to logging features that influenced seed dispersal patterns by comparing the mother–offspring distances for seedlings located inside versus outside canopy openings. Distance distributions were compared between groups (inside vs. outside openings) using a Wilcoxon rank-sum test (Wilcoxon, 1945), allowing for non-parametric comparison of dispersal ranges.

To investigate how logging disturbances affect genetic structure and diversity, we analyzed the following diversity metrics using SPAGeDi 1.5a (Hardy & Vekemans, 2002) for seedlings located inside and outside canopy openings: observed heterozygosity (*H*_O_), expected heterozygosity (*H*_E_), inbreeding coefficient (*F*_IS_), allelic richness (*A*_R_, standardized to 30 gene copies), selfing rate (s) and spatial genetic structure (*Sp*), hypothesising that seedlings in canopy openings may have lower genetic diversity, be more inbred and more related to each other at short distance than those outside such gaps. The significance of *F*_IS_ was evaluated through 10,000 permutations of alleles among individuals. The Sp statistic for nuclear and plastid markers was calculated following (Vekemans & Hardy, 2004), based on the slope of the regression of pairwise kinship coefficients (*F*_ij_) on the logarithm of spatial distance. Kinship-distance curves were generated to assess how genetic similarity decays with distance in gap vs. non-gap areas. The significance of the variations observed between the diversity indices (*A*_R_, *H*_E_, *H*_O_, *F*_IS_, Sp, s) was tested between plots and cohorts by bootstrapping (n= 10,000).

## 3. Results

### 3.1. Sample quality and clonality

Across the combined dataset of 703 individuals from the two study plots, multilocus genotype (MLG) screening identified two individuals with identical genotypes. Given their consecutive sampling identifiers and very close geographic coordinates, these were interpreted as a duplicate sampling of the same tree, and one was removed. A multilocus lineage (MLL) analysis with a conservative threshold (α² = 5) identified six additional cases of near-identical genotypes. Two of these corresponded to neighbouring wells in genotyping plates and were considered as contamination. These genotypes were removed, leaving a final dataset of 700 individuals.

Genotypic diversity estimated with RClone remained extremely high. The Shannon index reached H′ = 6.55 for MLGs and 6.54 for MLLs, i.e. Hill numbers of order one (¹D = exp(H′)) of 699 and 693 respectively, values close to the observed numbers of distinct multilocus genotypes (699) and lineages (695). This indicates that nearly all individuals carried a unique multilocus profile and that these profiles were evenly represented, i.e. a near-absence of clonality or genotypic redundancy in the dataset. Null allele analysis confirmed high genotyping quality, with mean frequencies of 0.010 ± 0.024 (HKO50) and 0.009 ± 0.022 (PAI74), and only 9-11% of markers exceeding the 0.05 threshold (Supporting information, Fig. S1). These results demonstrate the robustness of the genotyping, the very low occurrence of contamination or duplication, and the near-absence of clonality in *D. guianensis*, ensuring reliability for downstream demographic and genetic analyses.

### 3.2. Population structure

Pairwise F*_ST_* estimates computed across all loci revealed a low but significant level of genetic differentiation between the two study plots HKO50 and PAI74 (*F*_ST_ = 0.0137, P < 0.001). Bayesian clustering analyses conducted with STRUCTURE revealed high levels of admixture across individuals from both forest plots (Supporting information, Fig. S2). Although the ΔK method identified K = 2 as the most likely number of clusters, the likelihood estimates LnP(K) increased steadily as K increased, and thus did not allow for a reliable inference of the best solution for K (Supporting information, Fig. S3). When examining admixture patterns among plots and across cohorts, PAI74 showed a greater heterogeneity than HKO50, especially in the JUV group, which presented high admixture levels regardless of K value.

When the two plots were analyzed jointly, the first sPCA axis had individuals from the two sites distributed along it, emphasizing a gradual allele frequency gradient across the full study area consistent with STRUCTURE results (Supporting information, Fig. S4). The interpolation of lagged scores showed a gradual transition between the HKO50 and southeastern PAI74 zones (Supporting information, Fig. S5). Analyzed separately, both HKO50 and PAI74 exhibited significant global genetic structure (p < 0.0001), indicating that individuals located near each other were genetically more similar than expected by chance. The spatial distribution of the lagged scores within each plot (Supporting information, Fig.S5B, and Fig.S5C) revealed subtle gradients in genetic composition. Weak local structure was detected in PAI74 (p < 0.01), whereas no significant local structure was found in HKO50. PCA also revealed a genetic separation between plots with some overlap and no clear pattern was observed between cohorts (Supporting information, Fig.S6).

### 3.3. Plot and cohort genetic diversity and SGS

At the full plot level, there are no significant differences between the genetic metrics. Patterns of spatial genetic structure, quantified using the *Sp* statistic, were 15% stronger in HKO50 (*Sp* = 0.019) than in PAI74 (*Sp* = 0.016 for both time points), visually confirmed by the spatial autocorrelograms of the kinship coefficient (Figure 2, Supporting information Table S1).

**Figure 2.**
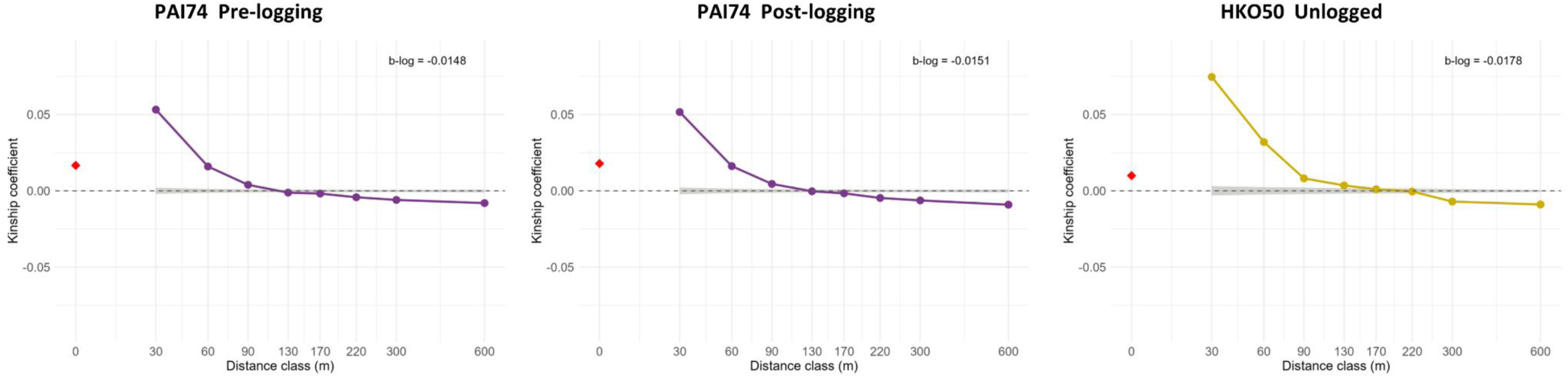
spatial autocorrelograms of the kinship coefficient across logged and unlogged conditions in PAI74 and HKO50. Kinship-distance curves (Loiselle’s kinship coefficient) for the full population of *D. guianensis* in the pre-logging PAI74, the post-logging PAI74, and the unlogged plot HKO50. Each point represents the mean pairwise kinship coefficient within a distance class, with the 0 m class (intra-individual) represented by a red rhombus. Shaded areas show the 95% confidence interval under the null model of random spatial distribution (10 000 permutations). Note that the 95% confidence intervals are particularly narrow, reflecting the high statistical precision of the estimates driven by the large sample sizes and high number of markers. The slope of the regression of kinship coefficients on the logarithm of distance (b-log) is indicated in each panel.

Analyzed independently, the different DBH-based cohorts (JUV, INT, ADL) revealed some contrasts in genetic diversity and spatial genetic structure between the two plots (Table 1). A significant difference was only observed between the two adult cohorts, with 77% of variation in selfing and 25% for the Sp statistic (Supporting information Table S1). Juveniles in PAI74 showed a significant inbreeding coefficient (*F*_IS_ = 0.023, p < 0.001) and the highest selfing rate (*S* = 0.031). In HKO50, *F*_IS_ in juveniles was also positive but weaker (*F*_IS_ = 0.017, p < 0.05), suggesting moderate inbreeding in both plots. At PAI74, intermediate cohorts showed higher private allelic richness than juveniles (0.383 vs. 0.279, variation of 37%), suggesting temporal structure in successive cohorts, whereas in HKO50 values were similar across cohorts (≈0.28-0.30). In HKO50, *F*_IS_ values were significantly lower in intermediate compared to juveniles and adults (respectively -0.041 and -0.033). The only significant selfing difference occurred between adult cohorts of PAI74 and HKO50, with lower selfing in logged adults (p<0.05) (Supporting information Table S1). In HKO50, Sp decreased from ADL (0.028) to INT (0.018), indicating weaker spatial genetic structure in the intermediate cohort compared to adults, while in PAI74, *Sp* remained mostly stable across cohorts (≈0.016–0.019). These trends were also visually supported by the kinship-distance correlograms (Supporting information, Fig. S7).

**Table 1.** Genetic diversity and spatial genetic structure for *D. guianensis* across the two forest plots and their cohorts. A_R_ (k=30): allelic richness standardized to 30 gene copies; pA_R_ (G=30): private allelic richness standardized to 30 gene copies; *H*_E_: expected heterozygosity; *H*_O_: observed heterozygosity; *F*_IS_: inbreeding coefficient; *F*_1_: mean pairwise kinship coefficient within the first distance class; *Sp*: intensity of spatial genetic structure, calculated from the regression of *F*_IJ_ on the logarithm of geographic distance; *NS*: Wright’s neighborhood size (1/*Sp*); *S*: selfing rate estimated from multilocus genotypes. Values significance : *, *P* < 0.05; **, P < 0.01; ***, *P* < 0.001; n.s.: not significant.

| Plots | Category | Sample size | $A_R(k=30)$ | $pA_R(G=30)$ | $H_E$ | $H_O$ | $F_{IS}$ | $F_1$ | $Sp$ | $NS$ | $S$ |
| --- | --- | --- | --- | --- | --- | --- | --- | --- | --- | --- | --- |
| PAI74 | All_unlogged | 417 | 4.12 | 0.175 | 0.552 | 0.541 | 0.019*** | 0.053 | 0.016*** | 62.5 | 0.020*** |
| PAI74 | All post logging | 375 | 4.13 | 0.180 | 0.552 | 0.541 | 0.020*** | 0.067 | 0.016*** | 62.5 | 0.022*** |
| PAI74 | ADL pre-logging | 135 | 4.10 | 0.274 | 0.549 | 0.540 | 0.015* | 0.066 | 0.019*** | 52.6 | 0.004 n.s. |
| PAI74 | ADL post-logging | 93 | 4.14 | 0.289 | 0.550 | 0.540 | 0.019* | 0.075 | 0.021*** | 47.6 | 0.006 n.s. |
| PAI74 | INT | 61 | 4.25 | 0.383 | 0.556 | 0.554 | 0.003 n.s. | 0.032 | 0.016*** | 62.5 | 0.011 n.s. |
| PAI74 | JUV | 221 | 4.07 | 0.279 | 0.551 | 0.538 | 0.023*** | 0.055 | 0.017*** | 58.8 | 0.031*** |
| HKO50 | All unlogged | 283 | 4.07 | 0.192 | 0.546 | 0.540 | 0.010 n.s. | 0.075 | 0.019*** | 52.6 | 0.014*** |
| HKO50 | ADL | 81 | 3.98 | 0.231 | 0.545 | 0.540 | 0.009 n.s. | 0.087 | 0.028*** | 35.7 | 0.026*** |
| HKO50 | INT | 45 | 4.10 | 0.280 | 0.546 | 0.559 | -0.024 n.s. | 0.058 | 0.018*** | 55.6 | 0.009 n.s. |
| HKO50 | JUV | 157 | 4.07 | 0.295 | 0.544 | 0.535 | 0.017* | 0.079 | 0.022*** | 45.5 | 0.008 n.s. |

### 3.4. Parentage analysis and dispersal dynamics

Parentage analyses revealed differences in seed and pollen dispersal distances between the two forest plots. The joint analysis (with congruence of CERVUS and COLONY analyses) recovered 184 offspring (38% of the 484 offspring tested) with both parents identified, 59 in HKO50 and 125 in PAI74. An additional 80 offspring (16.5%) had only one parent identified while the second seed or pollen parent remained unknown. A total of 220 offspring (45.5%) had no compatible sampled parent for either vector and corresponded most likely to parentage from outside the studied plots. COLONY identified 15 sibship clusters across the two plots. Cluster 1 grouped 239 offspring (JUV and INT) with at least one identified parent, predominantly originating from PAI74, while Cluster 3 comprised 222 offspring, mainly from HKO50 (Figure 3). For within-plot dispersal events, in HKO50, the mean seed dispersal distance was estimated to 32 m, and the mean pollen dispersal distance to 291 m. In contrast, within-plot seed and pollen dispersal distances in the selectively logged plot PAI74 were shorter on average: 25 m for seed and 148 m for pollen (Supporting information, Fig. S8). Wilcoxon rank sum tests revealed no significant differences between plots in terms of the number of offspring per parent or the seed dispersal distances of mothers. Maternal contributions and the gene flow from seeds remained broadly similar from one plot to another. In contrast, a significant difference (p < 0.05) was detected for pollen dispersal distances, which were higher for fathers of offspring located in HKO50 than for those of offspring located in PAI74 (Supporting information, Fig. S9, S10).

**Figure 3.**
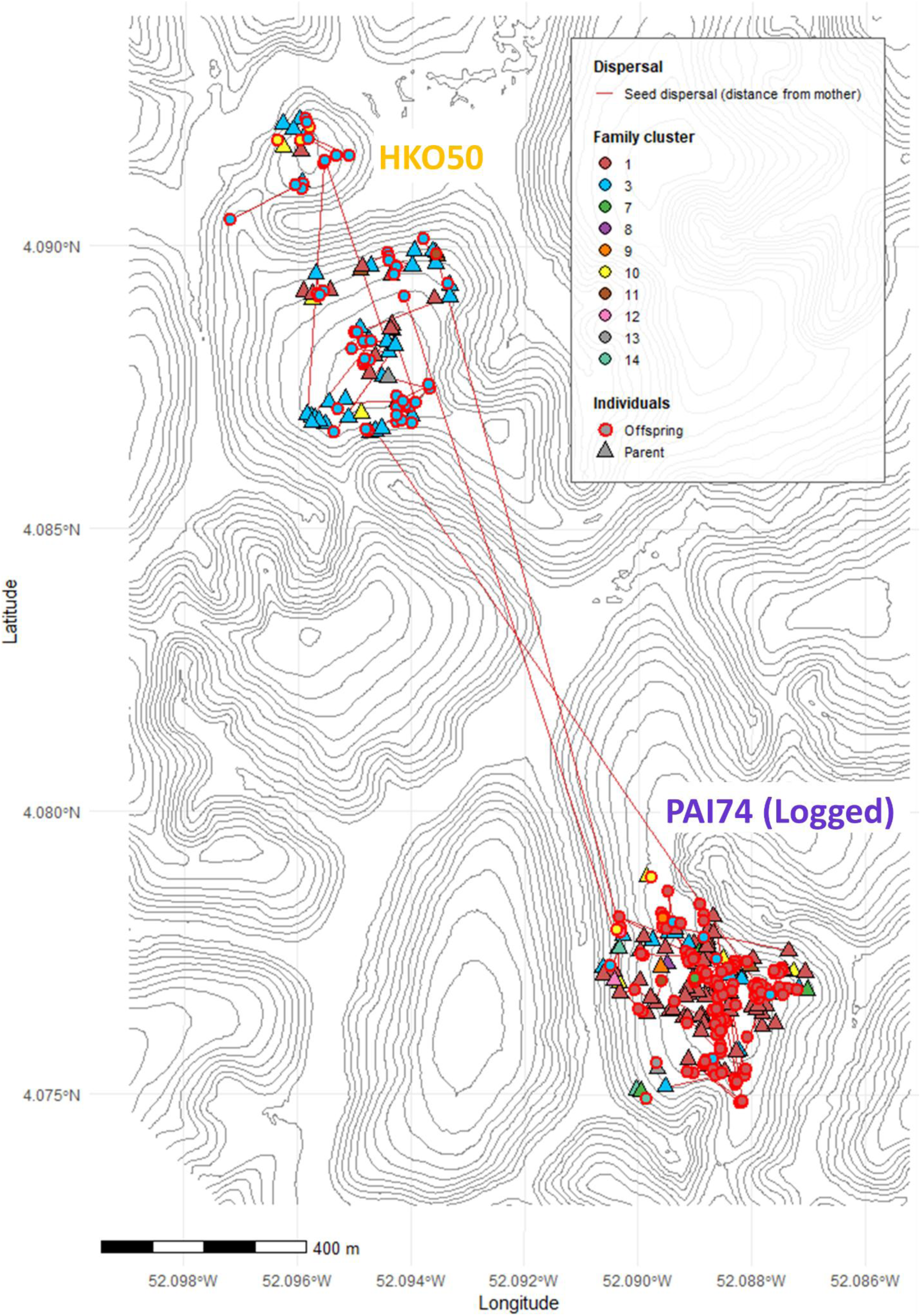
Spatial representation of mother–offspring dispersal events and family cluster inferred by COLONY. Maps showing seed dispersal events inferred from consistent parentage assignments (COLONY and CERVUS combined). Individuals are colored by family clusters based on full- and half-sib relationships detected using COLONY. Topographic isolines (5 m elevation intervals) illustrate the relief across both plots.

Parentage assignments also revealed differences in the direction and frequency of dispersal events (Figure 4). These patterns are based exclusively on the 184 offspring with both parents successfully identified (59 in HKO50, 125 in PAI74). Parent locations are reported by plot (59 parents in HKO50, 125 in PAI74), whereas offspring counts follow the plot in which the offspring was established (41 in HKO50, 143 in PAI74). Of the 184 offspring with an identified father (41 in HKO50, 143 in PAI74), most had the father located within the same plot: 30 out of 41 HKO50 offspring (73.2%) and 114 out of 143 PAI74 offspring (79.7%). Cross-plot pollen flow was nonetheless substantial, with 11 HKO50 offspring (26.8%) having a father originating from PAI74, and 29 PAI74 offspring (20.3%) having a father from HKO50, indicating active pollen exchange between the two plots in both directions. In contrast, seed dispersal was almost exclusively restricted to within plots: 58 out of 59 HKO50 offspring (98.3%) and 124 out of 125 PAI74 offspring (99.2%) had their mother located within the same plot.

**Figure 4.**
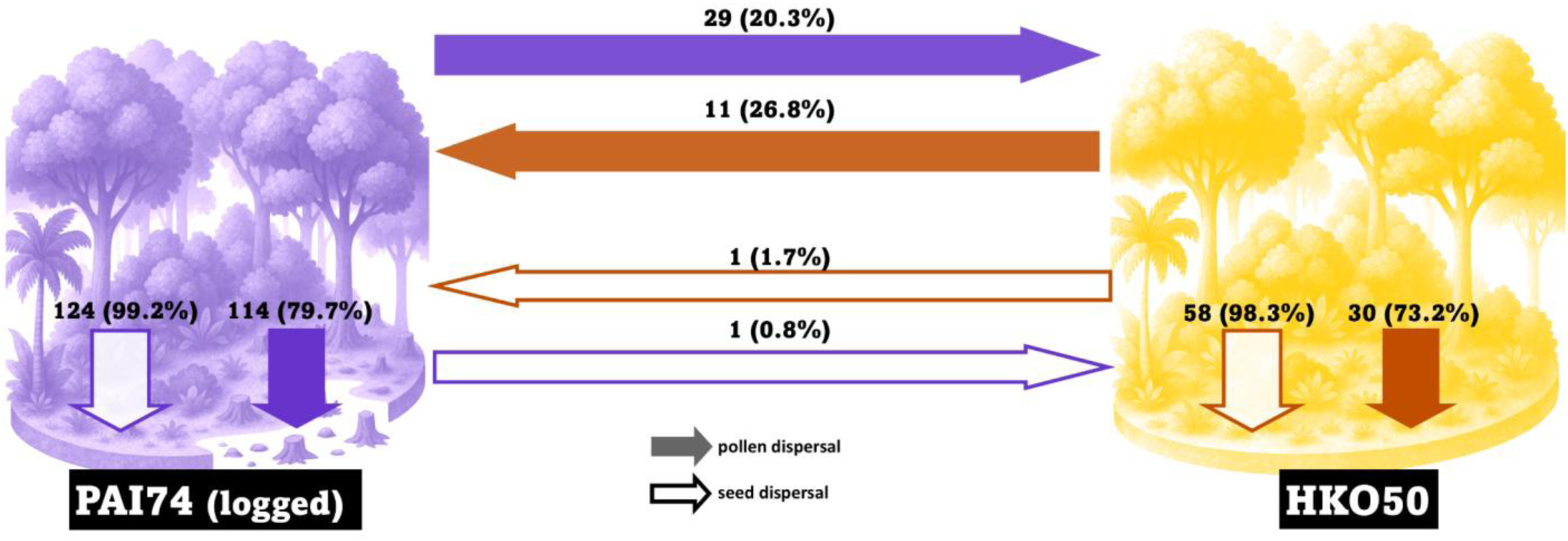
Graphical representation of seed and pollen dispersal events within and between plots. The arrows show the number of dispersal events and the associated percentage per plot (percentage per plot based on the 184 offsprings identified with two parents).

Among mothers, the distribution of reproductive success appeared relatively even across both plots, with similar Gini values (HKO50 = 0.41, PAI74 = 0.45). In contrast, among fathers we observed a balanced reproductive contribution in HKO50 (Gini = 0.29), but in PAI74 reproductive success was skewed (Gini = 0.47). Bootstrap comparisons of Gini indices did not yield statistically significant differences between plots (Supporting information, Fig. S11). The effective number of mothers was broadly similar between plots (HKO50: Ne_m = 15.6; PAI74: Ne_m = 26.0). Paternal effective numbers differed markedly between plots: in HKO50, Ne_f = 30.9, whereas in PAI74, Ne_f = 12.2. Among mothers, reproductive success showed a significant interaction with DBH only in PAI74 (p < 0.05). Maternal dispersal distances were not significantly related to DBH or plot, indicating that female tree size does not determine how far seeds are dispersed. Among fathers, neither reproductive success nor mean pollen dispersal distance was associated with DBH, and no differences emerged between plots.

Dispersal kernel estimation using NMπ2 confirmed the pattern of more restricted gene flow in PAI74 with median seed dispersal distance was estimated at 41.3 m v.s. 48.8 m in HKO50, and median pollen dispersal distance at 175.9 m in PAI74 versus 184.5 m in HKO50. Both plots showed high pollen immigration rates (PAI74: mp = 0.693; HKO50: mp = 0.754), indicating that the majority of pollen originated from outside the sampled plots, and substantial seed immigration rates (PAI74: ms = 0.378; HKO50: ms = 0.439). Taken together, these kernel-based estimates indicate slightly more restricted dispersal and lower immigration in the logged plot PAI74.

### 3.5. Post-logging regeneration dynamics

Spatial intersection analysis revealed a non-random seedling distribution in relation to logging-induced canopy features. A Firth logistic regression model confirmed the effect of these microhabitat disturbances on seedling occurrence (Likelihood ratio test = 286.47, df = 3, p < 0.0001), with a significantly higher proportion of seedlings found within felling gaps, skid trails, or log landings (Figure 5). The dispersal distances of seedlings were not influenced by disturbances. The distances between mothers and their offspring are similar for seedlings established inside canopy openings and those located outside.

**Figure 5.**
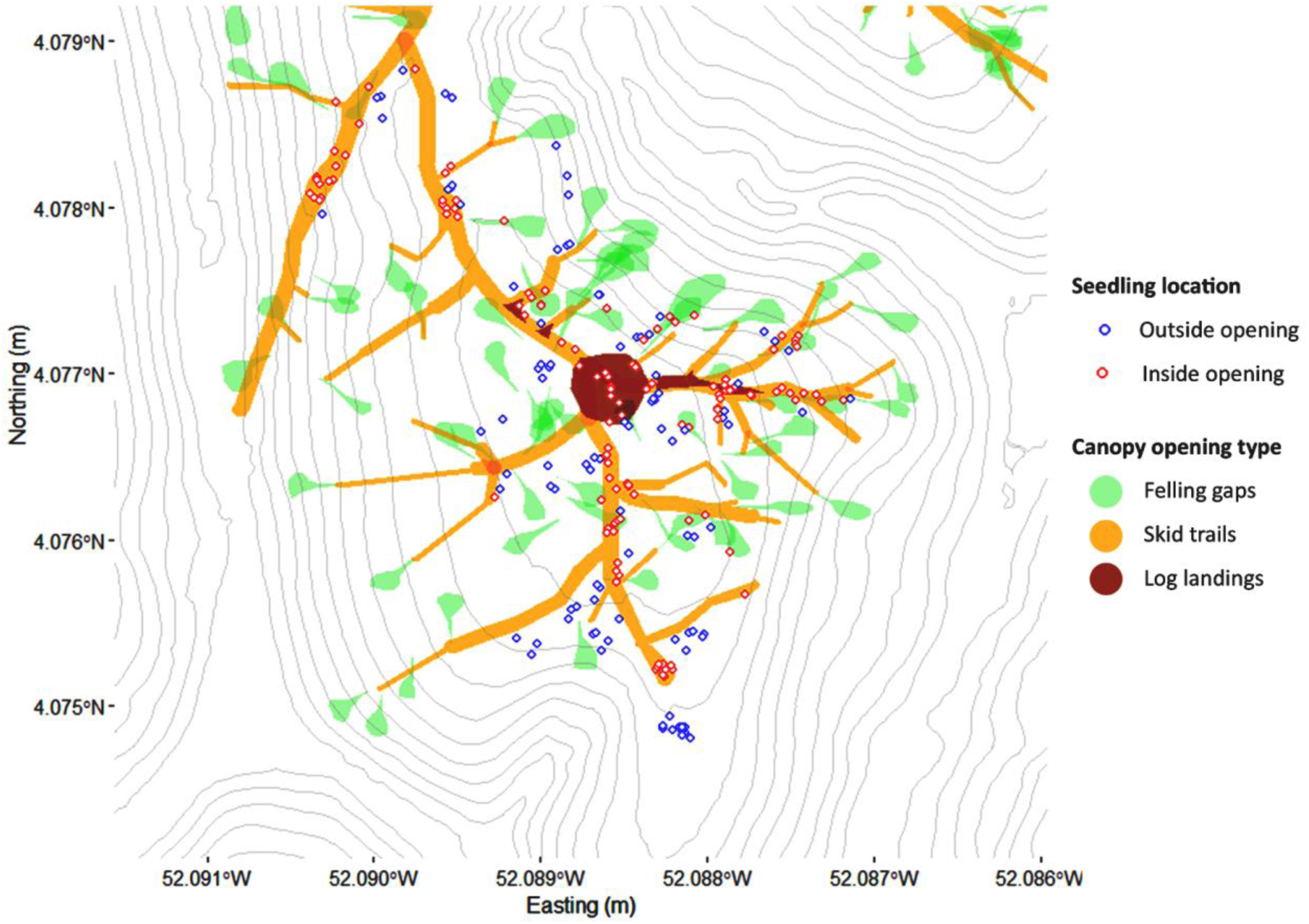
Spatial distribution of seedling establishment in relation to logging-induced canopy openings in plot PAI74. Map showing the location of seedlings (colored by canopy status) relative to three types of logging-related features: felling gaps (light green), skid trails (orange), and log landings (brown). Seedlings located inside disturbed areas are marked in red, while those outsides are shown in blue. Topographic isolines (5 m elevation intervals) illustrate the relief across both plots.

Genetic diversity metrics of seedlings established inside and outside canopy openings were broadly similar and not significantly different, with similar allelic richness (*A*_R_ = 4.04–4.08) and expected heterozygosity (*H*_E_ = 0.55) between micro-habitats (Table 2). Inbreeding levels (F*_IS_*) and selfing rates (S) remained low and did not differ significantly. In contrast, spatial genetic structure analyses revealed divergence between nuclear and chloroplast markers. Nuclear *Sp* values were low and similar between groups (*Sp* = 0.021 inside vs. 0.015 outside), reflecting the combined effects of seed and pollen flow. In contrast, chloroplast markers, which trace only seed-mediated gene flow, showed contrasting levels of structuring: seedlings outside gaps exhibited stronger structuring (*Sp* = 0.215) than those inside openings (*Sp* = 0.113), suggesting more mixed-origin recruitment within gaps than under a closed canopy.

**Table 2.**
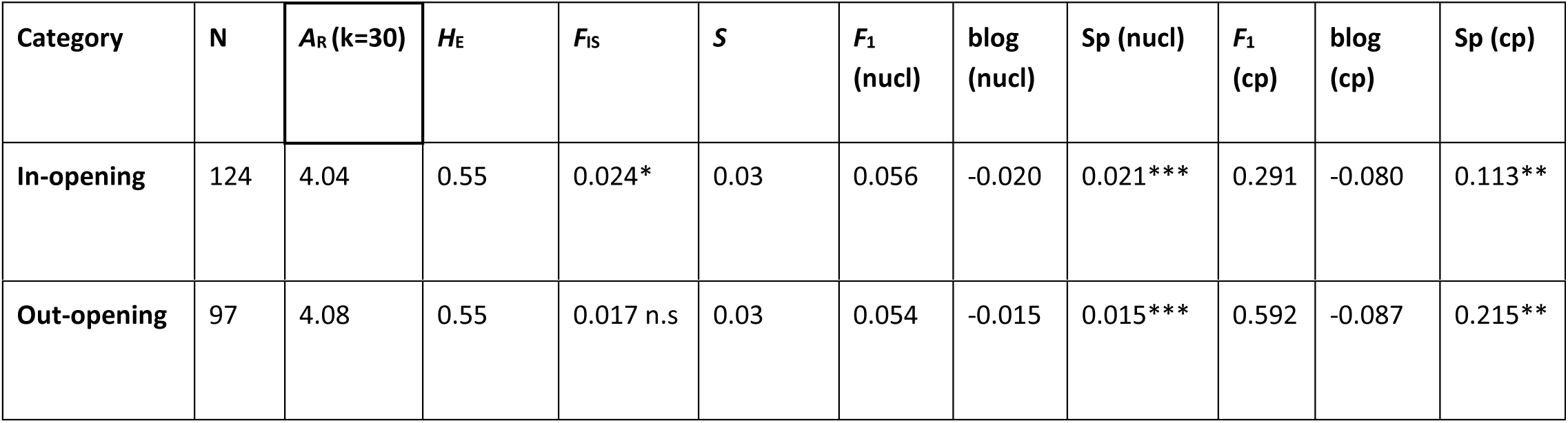
Genetic diversity and spatial genetic structure metrics for seedlings located inside versus outside logging-induced canopy openings in the PAI74 plot. *A*_R_(k = 30), allelic richness standardized to 30 gene copies; *H*_E_, expected heterozygosity; *S*, selfing rate estimated from heterozygote deficiency; *F*_IS_, inbreeding coefficient; *Sp*, spatial genetic structure intensity; *F*₁, average pairwise kinship coefficient in the first distance class; b-log, slope of the regression of kinship on log(distance); Sp, *F*₁ and b-log for both nuclear and plastid markers ; significance of value: *, P < 0.05;**, P < 0.01;***, P < 0.001;n.s., not significant.

## 4. Discussion

### 4.1. Genetic structuring and connectivity between plots

Despite the short geographic distance between the two study plots (∼1.5 km apart), our analyses revealed a weak but significant genetic differentiation (*F*_ST_ = 0.0137), consistent with patterns commonly observed in tropical trees where dispersal is spatially restricted (Dick et al., 2008; Hardy et al., 2006). The extensive admixture observed in STRUCTURE barplots (Supporting information, Fig.S2, S3), the apparent signal of K = 2 likely reflects isolation by distance rather than a true subdivision into distinct populations (see also Frantz et al., 2009). The strength of our microsatellite dataset allowed us to detect this very fine-scale differentiation, which likely results from geographic distance alone rather than from any barrier to gene flow. STRUCTURE analyses and sPCA both suggested allele frequency gradients reminiscent of isolation by distance (Supporting information, Fig S2, S3, S4), supporting the view that individuals from the two sites form a single, connected population. The two plots do not exhibit marked genetic structuration that would indicate distinct population subdivisions. The genetic differentiation observed among individuals reflects isolation-by-distance at the plot scale, which our high-resolution dataset was sufficiently powerful to detect even at fine spatial scales. Overall, our results suggest that gene flow is not absent nor strongly limited, but rather sufficient to maintain genetic cohesion across the two plots.

STRUCTURE results also revealed subtle within-plot heterogeneity, particularly in the PAI74 seedling cohort, which exhibited elevated admixture levels across a range of K values. This suggests that recent recruitment may integrate genetic input from multiple reproductive sources permitted by long distance pollen flow (Arruda et al., 2015; Born et al., 2008) (Supporting information, Fig.S10). This could also reflect the heterogeneity induced by logging in microhabitats already known to increase functional and species diversity in rainforest logged areas (Baraloto et al., 2012). HKO50 presented more homogeneous reproductive cluster assignments, especially among adult and intermediate trees, consistent with long-term demographic stability in the absence of disturbance. The weak local structure only observed in the PAI74 plot may be a first sign suggesting logging may have accentuated genetic clustering at fine spatial scales.

### 4.2. Variations in genetic diversity and fine scale structure

Despite contrasting management histories, genetic diversity metrics were overall similar between the unlogged plot HKO50 and the selectively logged plot PAI74 (Supporting information, Fig. S8). Allelic richness, expected and observed heterozygosity showed only minor differences between plots, and private allelic richness remained stable across management regimes. This stability contrasts with expectations that logging may erode genetic diversity, as observed in several tropical tree species such as *Iriartea deltoidea* (Sezen et al., 2005, 2007), *Jacaranda copaia* (Leclerc et al., 2015), and *Carapa guianensis* (André et al., 2008). However, our findings are consistent with other studies reporting no significant genetic erosion following logging, particularly when gene flow compensates for the demographic disturbance (Cloutier et al., 2007; Degen et al., 2006; Silva et al., 2008).

At both plots, the intermediate cohort showed slightly higher allelic richness compared with adults, although the magnitude of this difference was modest (Table 1, Supporting information Table S1). While this may reflect subtle demographic variation following logging, the differences are too small to infer strong shifts in genetic diversity. In PAI74, inbreeding coefficients and selfing rates in seedlings (*F*_IS_ = 0.023, *S* = 0.031) were marginally higher than in other cohorts, but again the variation was limited. In contrast, the unlogged plot HKO50 displayed broadly stable diversity indices across cohorts, with a slight increase in SGS from intermediate (Sp = 0.018) to adults (Sp = 0.028), consistent with the fine-scale kin structure typically reported for *D. guianensis*.

Overall, differences across cohorts in both plots were subtle, suggesting that genetic diversity has remained largely stable under both logged and unlogged conditions and across cohorts. Genetic metrics seem more strongly driven by life-stage dynamics than by logging history. No strong differences were observed between post-logging, pre-logging and unlogged adult trees (Supporting information, Table S1). Showing that the first logging cycle on PAI74 does not appear to have had a major impact on adult population diversity in contrast to the results suggested by Wernsdörfer et al. (2011) on *D. guianensis* by modelling logging impact.

### 4.3. Seed and pollen dispersal dynamics

Across both study plots, seed and pollen dispersal distances in *D. guianensis* were consistent with previous reports for the species in French Guiana (Bonnier et al., 2026; Latouche-Hallé et al., 2004). Mean dispersal distances were shorter in the logged plot (PAI74) compared to the unlogged plot (HKO50), for both seed (25 m vs. 32 m) and pollen (148 m vs. 291 m). Following NMπ2 results, the slightly lower seed and pollen dispersal distances, combined with reduced immigration rates in PAI74 compared to HKO50, are consistent with a moderate contraction of gene flow following selective logging. However, given the persistence of high immigration rates in the logged plot, this contraction does not appear severe enough to compromise genetic diversity in the short term, supporting the overall conclusion of resilience of *D. guianensis* to reduced-impact logging. Reductions in dispersal following disturbance have been documented in other tropical tree species, where logging promotes localized seed aggregation near maternal trees, thereby decreasing dispersal distances and strengthening fine-scale genetic clustering (Angbonda et al., 2021; Hardy et al., 2019). This effect may be especially pronounced in *D. guianensis*, whose seeds are suspected to undergo secondary dispersal by animals, potentially parrots, as observed in preliminary field studies (Bonnier et al., 2026). Logging operations coupled with hunting can disrupt the seed and pollen dispersers activity, either through direct habitat disturbance or a reduction in resource availability (Jennings et al., 2001; Poulsen et al., 2011).

In contrast to these localized seed shadows, parentage assignments highlighted the occurrence of long-distance pollen-mediated gene flow, including important exchange between plots already observed in neotropical trees (Sujii et al., 2021). The high proportion of offspring with at least one unidentified parent also underscores the critical role of long distance gene dispersal in sustaining genetic diversity and connectivity in *D. guianensis*. This pattern was observed in both the logged and unlogged plot and consistent with the NMπ2 results, suggesting that reliance on external gene flow is an intrinsic feature of the reproductive system of *D. guianensis*. These results are supported by the sibship cluster assignment from COLONY, showing that individuals from the two main clusters are distributed between the two plots, showing a strong connectivity. These results corroborate earlier studies showing that pollen flow often exceeds seed dispersal distances and represents a dominant component of gene flow in trees (Latouche-Hallé et al., 2004; Petit & Hampe, 2006). Long-distance connectivity is critical in perturbed landscapes by selective logging or fragmentation, where reduced adult density can constrain local mate availability (Wang et al., 2011). Indeed, reductions in conspecific density have been shown to increase pollinator foraging ranges, thereby extending effective pollen dispersal distances (Duminil, Daïnou, et al., 2016). This mechanism may buffer against the potential loss of genetic diversity by ensuring continued admixture, as also evidenced by the elevated levels of admixture in the seedling cohort of PAI74.

Reproductive success analyses revealed contrasts between maternal and paternal contributions. Among mothers, reproductive success was evenly distributed across plots but male reproductive success was skewed in PAI74 compared to a more balanced distribution in HKO50. Such reproductive skew reduces the effective number of breeders, increasing relatedness among seedlings and potentially accelerating inbreeding under certain conditions (Aldrich & Hamrick, 1998; Leclerc et al., 2015). The effective number of parents provided a standardized quantification of this reproductive skew. In *D. guianensis*, this asymmetry in favour of males may result from its ecological and reproductive characteristics: seedlings prefer to establish themselves in canopy openings and flowering is asynchronous depending on the year (Jésel, 2005). Individuals that flower in the same year as logging on the plot have a disproportionate advantage in colonising these canopy openings.

### 4.4. Impact of canopy openings in post logging regeneration

Selective logging operations in tropical forests inevitably open the canopy, leading to substantial changes in light, microclimate, and soil conditions that promote seedling establishment by reducing competition and enhancing resource availability (Clark & Covey, 2012; Putz et al., 2008; Swaine & Agyeman, 2008). Our results confirm this ecological facilitation for *D. guianensis*, with a significantly higher proportion of seedlings found within logging-induced microhabitats such as felling gaps. They are consistent with earlier observations of Jésel (2005) for *D. guianensis* and in other tropical species such as *Coula edulis*, where disturbed areas experienced better regeneration than intact forest, likely due to decreased interspecific competition and herbivory pressure (Kamdem et al., 2025). Canopy openings have been shown to stimulate fruiting in smaller trees due to enhanced light availability (Appanah & Manaf, 1990). This phenomenon has been observed in previous studies on *D. guianensis*, where higher seedling densities and survival rates were recorded in gaps compared to closed-canopy sites (Meer et al., 1998). Wind-dispersed species may benefit disproportionately from such microhabitats, as suggested by (Augspurger and Franson 1988), who reported enhanced seed rain into canopy gaps for anemochorous species. It is worth noting that, within logging-induced microhabitats, seedling establishment appeared to occur predominantly in skid trails rather than in felling gaps (Figure 5). This pattern likely reflects the combined effect of wider canopy openings and completely bare mineral soil in skid trails, which together create large, vegetation-free areas highly favorable for seedling establishment. In contrast, felling gaps tend to be narrower and retain a proportion of the surrounding understory vegetation after log extraction, resulting in reduced available space for new recruits.

Despite this demographic signal, genetic diversity metrics were broadly similar between seedlings inside and outside canopy gaps. Allelic richness (*A_R_*), expected heterozygosity (*H_E_*), inbreeding (*F_IS_*), and selfing rates (*S*) showed no significant differences, indicating that the genetic consequences of regeneration in canopy openings remain limited. The main contrast comes from spatial genetic structure (SGS). Nuclear markers revealed uniformly low and comparable structuring inside and outside gaps, and chloroplast markers, which trace only seed-mediated dispersal, showed significantly stronger structuring outside gaps. This suggests that seedlings recruited within gaps originate from a broader set of maternal sources, whereas recruitment outside gaps is more spatially restricted. Such a reduction of SGS inside canopy openings contrasts with findings in *J. copaia*, where gaps increased kin aggregation and SGS among seedlings (Leclerc et al., 2015). It should be noted that part of the juvenile cohort in PAI74 may have been established before logging. This “surviving” regeneration, facilitated by the opening of the canopy, could partially mask the genetic signature of post-logging recruitment, which would explain the absence of marked differences with the unlogged plot and juveniles out of the canopy opening.

It is important to stress that the apparent demographic success of seedlings in canopy openings may not translate into long-term recruitment. Jésel et al. (2005) showed that while canopy openings promote germination and early establishment, this advantage is transient: a decade after logging, juvenile recruitment declines and may fall below levels in undisturbed forest. *D. guianensis* regeneration trajectories are discontinuous, with height growth favored in gaps but later constrained by competition from heliophilous species. Multiple successive gap events are required for juveniles to reach the canopy. Our results confirm that logging-induced canopy openings create short-term opportunities for *D. guianensis* regeneration without immediate positive or negative impacts on genetic diversity. Given the discontinuous and competitive regeneration ecology of this species, the long-term demographic and genetic consequences remain uncertain and require evaluation beyond the first decade after logging.

### 4.5. Forest management implication

The persistence of high genetic diversity in adult trees and seedlings suggests that the current management regime succeeded in avoiding strong genetic erosion. However, early signs of altered dispersal dynamics, reproductive skew, and regeneration patterns in logged plots indicate that longer-term risks may emerge if logging is repeated without safeguards.

Regeneration of *D. guianensis* is associated with logging-induced canopy openings, including gaps and skid trails. While these disturbed microhabitats promote recruitment in the short term by increasing light and reducing competition, their demographic value is uncertain. Skid trails are likely to be reused in future harvests, destroying regeneration hotspots established after the first cycle. It therefore seems important in the following logging cycles to preserve those hotspots that will not be reused, such as felling gaps and some skid trails.

Our findings emphasize the buffering role of long-distance pollen flow. While seed dispersal remained mainly short-ranged and localized, pollen-mediated connectivity between logged and unlogged stands was substantial. Maintaining adjacent unlogged plots or lightly disturbed stands is therefore critical to sustaining gene flow and regeneration potential.

An important concern in selective logging practices is whether the removal of the largest, most phenotypically developed individuals constitutes dysgenic selection, potentially compromising the genetic quality of future generations. Studies across tropical forests have demonstrated that such practices can lead to changes in genetic metrics in regenerating populations (Degen et al., 2006; Johns, 1988; Sebbenn et al., 2008). While our study cannot directly assess the genetic “quality” of logged versus retained trees, the maintenance of genetic diversity and reproductive success across different tree sizes in the logged plot suggests that the remaining gene pool retains adequate genetic potential for population persistence over the 12-year timeframe examined.

*D. guianensis* seedlings establish successfully in canopy openings created by logging, but the sustainability of this regeneration remains uncertain beyond the first decade. Previous studies have shown that the demographic advantage conferred by gaps can rapidly decline after 10–15 years as competition from heliophilous species intensifies (Jésel, 2005). Like for fragmentation studies on forest trees, the impact of these disturbances could emerge during future generations (Lowe et al., 2005; Vranckx et al., 2012). To capture these delayed dynamics, we recommend implementing systematic genetic monitoring of logged stands at intervals of 15–20 years, extending throughout the entire 65-year cutting cycle. Such follow-up would make it possible to track the survival genetic diversity of regeneration cohorts over time, to evaluate whether the initial boost in seedling establishment provided by canopy openings translates into long-term population renewal.

## 5. Conclusion

This study provides a comprehensive assessment of the demographic and genetic impacts of selective logging on *D. guianensis*. Overall genetic diversity and adult structure were not significantly altered by logging, suggesting short-term resilience of the reproductive population. Subtle shifts emerged in dispersal dynamics and reproductive success: seed dispersal remained localized and stable, whereas pollen dispersal was reduced in the logged plot and reproductive skew was more pronounced among fathers. Seedlings established preferentially in canopy openings, reflecting the facilitation of regeneration by gaps, yet genetic diversity and inbreeding levels remained broadly similar across microhabitats. Taken together, these results highlight the resilience of *D. guianensis* to low-intensity selective logging, while also emphasizing potential risks of altered reproductive dynamics. They underscore the importance of long-term monitoring to ensure that short-term regeneration success effectively translates into sustainable population renewal across harvest cycles.

## Declarations

### Authors’ contribution

Designed research: JB, OB, ST, NT, MH; performed research: JB, VT, NT, MH; performed the sampling: JB, NT; analyzed data: JB; wrote the paper: JB, NT, MH. All authors critically read and approved the submitted version of the paper.

## Acknowledgments

We are especially grateful to St-Omer Cazal for being a major help during the sampling and DNA extraction for this project. We also thank the master’s students in the Tropical Rainforest (FTH) module of 2024 for their help during a week of sampling at the HKO50 forest plot. We thank Enrique Saez-Laguna for his help during sampling in Régina. Microsatellite development and genotyping by sequencing were performed at the PGTB (https://doi.org/10.15454/1.5572396583599417E12) with the help of Olivier Lepais, Zoé Compagnie, Emilie Chancerel, Christophe Boury, and Erwan Guichoux.

## Availability of data

The datasets generated and analysed during the current study are available in the Zenodo repository, https://doi.org/10.5281/zenodo.18405085

## Ethics approval

The authors confirm that they follow the good scientific practice and all ethical standards requested by the journal. The authors have obtained access to genetic resources on the French national territory (APA standards, file number: 8360908).

## Consent for publication

All authors gave their informed consent to this publication and its content.

