## Supporting information 1 for "Impact of selective logging on the population dynamics and genetic diversity in the neotropical timber tree *Dicorynia guianensis*"

The following figures and tables provide additional results supporting the analyses presented in the main manuscript. These supplementary materials include detailed outputs, extended visualizations, and complementary statistical results that could not be presented in the main text for reasons of space. They are intended to facilitate transparency and reproducibility of the study by providing access to all relevant data and graphical outputs.

| **Type** | **Comparison** | **Variable** | **Percent_Diff** | **Signifiance** |
| --- | --- | --- | --- | --- |
| Inter-plot | HKO50 vs PAI74: All | *A*_R_ | -1.47 | n.s. |
| Inter-plot | HKO50 vs PAI74: JUV | *A*_R_ | 0 | n.s. |
| Inter-plot | HKO50 vs PAI74:: INT | *A*_R_ | -3.66 | n.s. |
| Inter-plot | HKO50 vs PAI74: ADL | *A*_R_ | -4.02 | n.s. |
| Intra-plot | PAI74: INT vs JUV | *A*_R_ | 4.42 | p<0.05 |
| Intra-plot | PAI74: ADL vs JUV | *A*_R_ | 1.72 | n.s. |
| Intra-plot | PAI74: ADL vs INT | *A*_R_ | -2.59 | p<0.05 |
| Intra-plot | HKO50: INT vs JUV | *A*_R_ | 0.74 | n.s. |
| Intra-plot | HKO50: ADL vs JUV | *A*_R_ | -2.21 | n.s. |
| Intra-plot | HKO50: ADL vs INT | *A*_R_ | -2.93 | p<0.05 |
| Intra-plot | PAI74: ADL post-logging vs ADL pre-logging | *A*_R_ | -0.97 | n.s. |
| Inter-plot | HKO50 vs PAI74: All | *F*_IS_ | -100 | n.s. |
| Inter-plot | HKO50 vs PAI74: JUV | *F*_IS_ | -35.29 | n.s. |
| Inter-plot | HKO50 vs PAI74: INT | *F*_IS_ | 112.5 | n.s. |
| Inter-plot | HKO50 vs PAI74: ADL | *F*_IS_ | -111.11 | n.s. |
| Intra-plot | PAI74: INT vs JUV | *F*_IS_ | -86.96 | n.s. |
| Intra-plot | PAI74: ADL vs JUV | *F*_IS_ | -17.39 | n.s. |
| Intra-plot | PAI74: ADL vs INT | *F*_IS_ | 533.33 | n.s. |
| Intra-plot | HKO50: INT vs JUV | *F_I_*_S_ | -241.18 | p<0.05 |
| Intra-plot | HKO50: ADL vs JUV | *F*_IS_ | -47.06 | n.s. |
| Intra-plot | HKO50: ADL vs INT | *F*_IS_ | -137.5 | p<0.05 |
| Intra-plot | PAI74: ADL post-logging vs ADL pre-logging | *F_I_*_S_ | -21.05 | n.s. |
| Inter-plot | HKO50 vs PAI74: All | *H*_E_ | -1.1 | n.s. |
| Inter-plot | HKO50 vs PAI74: JUV | *H*_E_ | -1.29 | n.s. |
| Inter-plot | HKO50 vs PAI74: INT | *H*_E_ | -1.83 | n.s. |
| Inter-plot | HKO50 vs PAI74: ADL | *H*_E_ | -0.92 | n.s. |
| Intra-plot | PAI74: INT vs JUV | *H*_E_ | 0.91 | n.s. |
| Intra-plot | PAI74: ADL vs JUV | *H*_E_ | -0.18 | n.s. |
| Intra-plot | PAI74: ADL vs INT | *H*_E_ | -1.08 | n.s. |
| Intra-plot | HKO50: INT vs JUV | *H*_E_ | 0.37 | n.s. |
| Intra-plot | HKO50: ADL vs JUV | *H*_E_ | 0.18 | n.s. |
| Intra-plot | HKO50: ADL vs INT | *H*_E_ | -0.18 | n.s. |
| Intra-plot | PAI74: ADL post-logging vs ADL pre-logging | *H*_E_ | -0.18 | n.s. |
| Inter-plot | HKO50 vs PAI74: All | *H*_O_ | -0.19 | n.s. |
| Inter-plot | HKO50 vs PAI74: JUV | *H*_O_ | -0.56 | n.s. |
| Inter-plot | HKO50 vs PAI74: INT | *H*_O_ | 0.89 | n.s. |
| Inter-plot | HKO50 vs PAI74: ADL | *H*_O_ | 0 | n.s. |
| Intra-plot | PAI74: INT vs JUV | *H*_O_ | 2.97 | n.s. |
| Intra-plot | PAI74: ADL vs JUV | *H*_O_ | 0.37 | n.s. |
| Intra-plot | PAI74: ADL vs INT | *H*_O_ | -2.53 | n.s. |
| Intra-plot | HKO50: INT vs JUV | *H*_O_ | 4.49 | p<0.05 |
| Intra-plot | HKO50: ADL vs JUV | *H*_O_ | 0.93 | n.s. |
| Intra-plot | HKO50: ADL vs INT | *H*_O_ | -3.4 | n.s. |
| Intra-plot | PAI74: ADL post-logging vs ADL pre-logging | *H*_O_ | 0 | n.s. |
| Inter-plot | HKO50 vs PAI74: JUV | *s* | -287.5 | n.s. |
| Inter-plot | HKO50 vs PAI74: INT | *s* | -22.22 | n.s. |
| Inter-plot | HKO50 vs PAI74: ADL | *s* | 76.92 | p<0.05 |
| Intra-plot | PAI74: INT vs JUV | *s* | -64.52 | n.s. |
| Intra-plot | PAI74: ADL vs JUV | *s* | -80.65 | n.s. |
| Intra-plot | PAI74: ADL vs INT | *s* | -45.45 | n.s. |
| Intra-plot | HKO50: INT vs JUV | *s* | 12.5 | n.s. |
| Intra-plot | HKO50: ADL vs JUV | *s* | 225 | n.s. |
| Intra-plot | HKO50: ADL vs INT | *s* | 188.89 | n.s. |
| Intra-plot | PAI74: ADL post-logging vs ADL pre-logging | *s* | -33.33 | n.s. |
| Inter-plot | HKO50 vs PAI74: JUV | *Sp* | 15.79 | n.s. |
| Inter-plot | HKO50 vs PAI74: INT | *Sp* | 22.73 | n.s. |
| Inter-plot | HKO50 vs PAI74: ADL | *Sp* | 11.11 | n.s. |
| Inter-plot | HKO50 vs PAI74: JUV | *Sp* | 25 | p<0.05 |
| Intra-plot | PAI74: INT vs JUV | *Sp* | -5.88 | n.s. |
| Intra-plot | PAI74: ADL vs JUV | *Sp* | 23.53 | n.s. |
| Intra-plot | PAI74: ADL vs INT | *Sp* | 31.25 | n.s. |
| Intra-plot | HKO50: INT vs JUV | *Sp* | -18.18 | n.s. |
| Intra-plot | HKO50: ADL vs JUV | *Sp* | 27.27 | n.s. |
| Intra-plot | HKO50: ADL vs INT | *Sp* | 55.56 | n.s. |
| Intra-plot | PAI74: ADL post-logging vs ADL pre-logging | *Sp* | -9.52 | n.s. |
| Inter-plot | HKO50 vs PAI74: JUV | *pA_R_* | 8.85 | n.t. |
| Inter-plot | HKO50 vs PAI74: INT | *pA_R_* | 5.42 | n.t. |
| Inter-plot | HKO50 vs PAI74: ADL | *pA_R_* | -36.79 | n.t. |
| Inter-plot | HKO50 vs PAI74: JUV | *pA_R_* | -25.11 | n.t. |
| Intra-plot | PAI74: INT vs JUV | *pA_R_* | 37.28 | n.t. |
| Intra-plot | PAI74: ADL vs JUV | *pA_R_* | 3.58 | n.t. |
| Intra-plot | PAI74: ADL vs INT | *pA_R_* | -24.54 | n.t. |
| Intra-plot | HKO50: INT vs JUV | *pA_R_* | -5.08 | n.t. |
| Intra-plot | HKO50: ADL vs JUV | *pA_R_* | -21.69 | n.t. |
| Intra-plot | HKO50: ADL vs INT | *pA_R_* | -17.50 | n.t. |
| Intra-plot | PAI74: ADL post-logging vs ADL pre-logging | *pA_R_* | 5.47 | n.t. |

**Table S1. Comparative differences in genetic diversity metrics between plots and cohorts of *Dicorynia guianensis*.** Percent differences (Percent_Diff) in allelic richness (*A*_R_), inbreeding coefficient (*F*_IS_), expected heterozygosity (*H*_E_), observed heterozygosity (*H*_O_), selfing rate (s) and spatial genetic structure (*Sp*) are shown for inter-plot comparisons (PAI74 vs. HKO50), intra-plot comparisons among cohorts (juveniles, intermediates, adults), and pre- vs. post-logging adults in PAI74. Significance was assessed by locus-based bootstrapping (n = 1000); n.s. : not significant ; n.t. not tested.

**Figure S1. Null allele frequency estimates for 66 nuclear SSR markers.** Brookfield estimates with 95% bootstrap confidence intervals for PAI74 (top) and HKO50 (bottom). Red bars indicate frequencies >0.05 (threshold shown by dashed line).
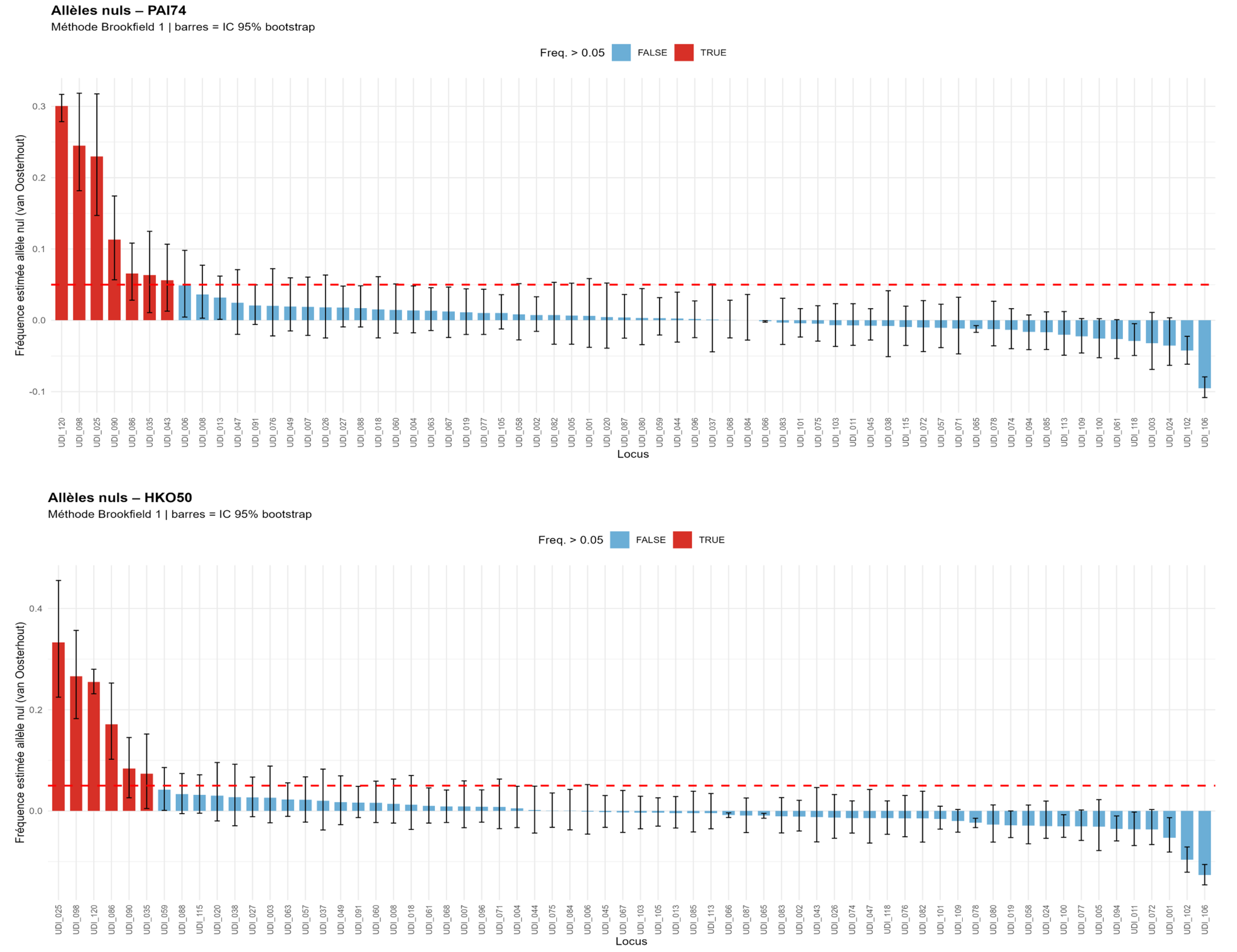

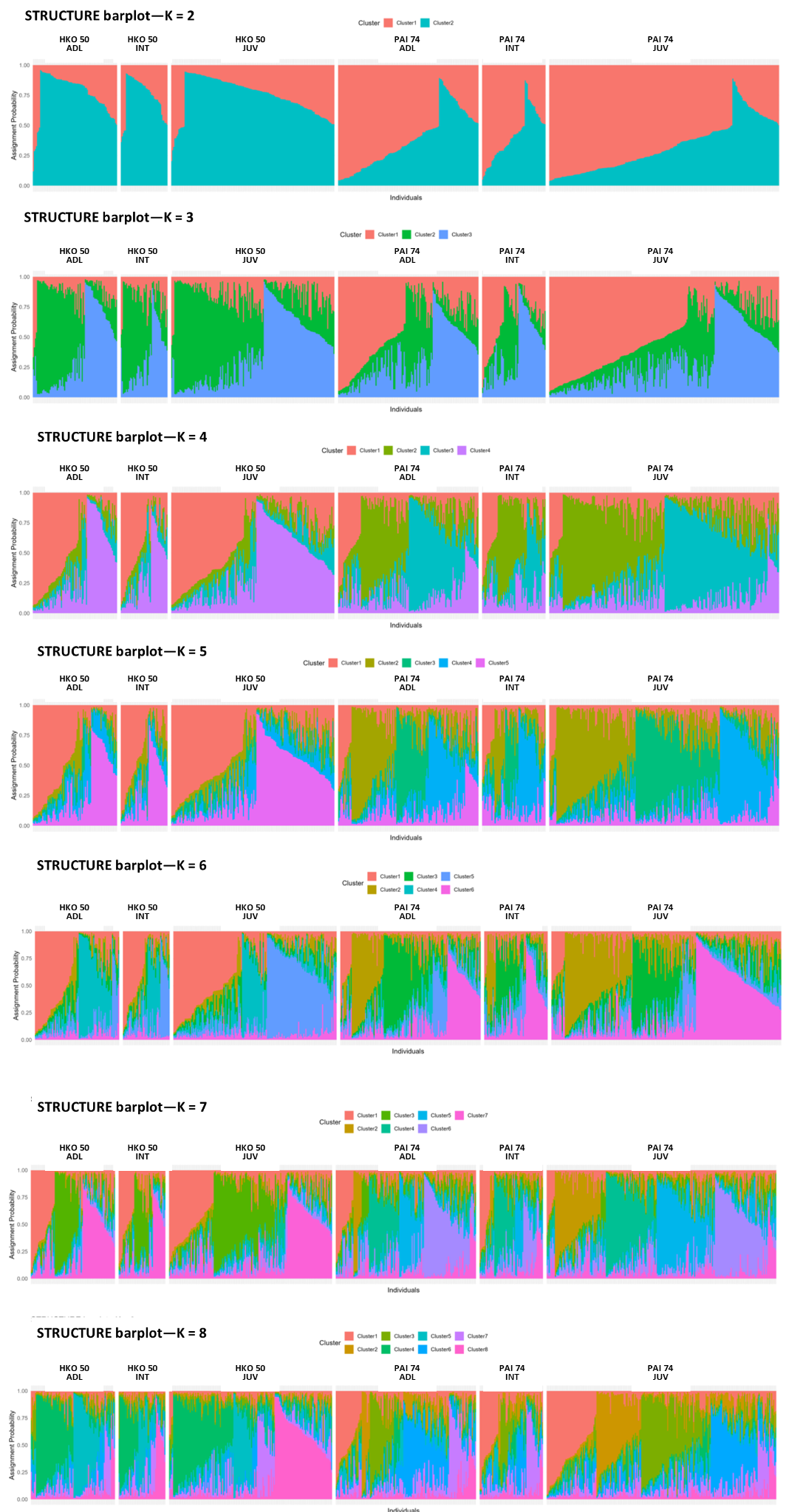

**Figure S2. Bayesian estimates of population structure based on nuclear SSR data in sampling sites, from K=2 to K=8.** Each vertical line represents one individual, and color segments represent its assignment proportions to each of K genetic clusters. Individuals are grouped by cohorts (SED, INT, ADL) and by plots (HKO50, PAI74).

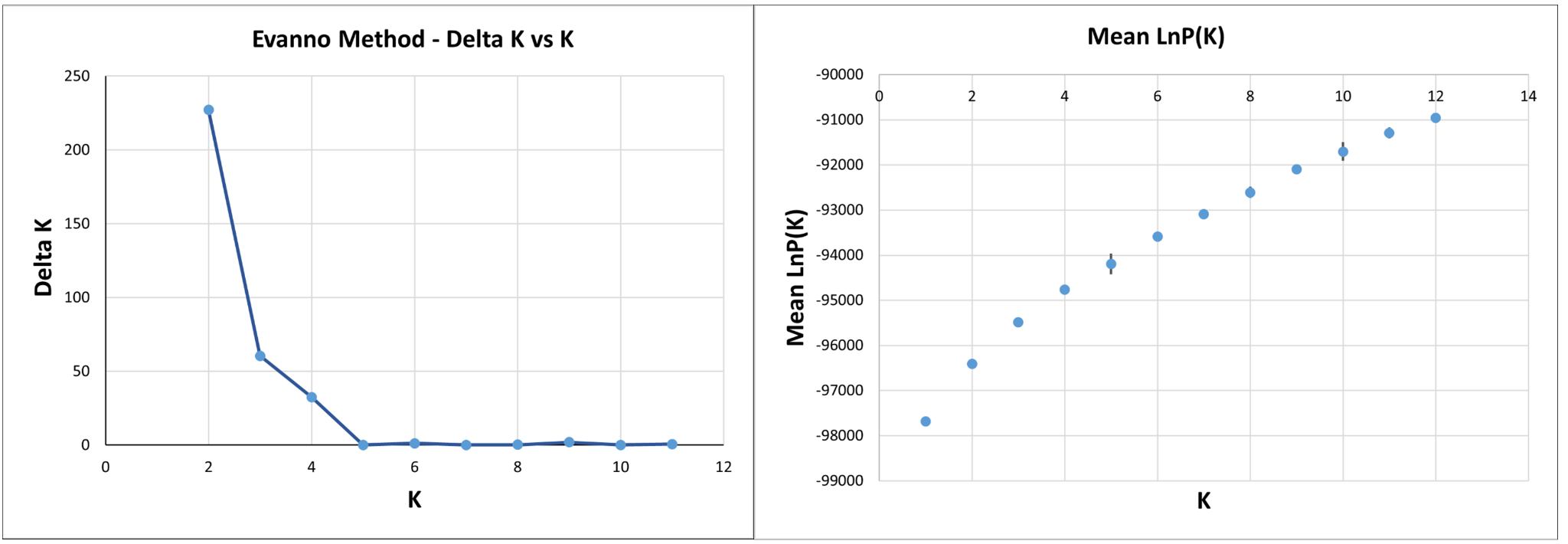

**Figure S3. Inference of the optimal number of genetic clusters (K) using ΔK and Mean LnP method from STRUCTURE analyses. A.** Delta K values from the mean log-likelihood probabilities from STRUCTURE runs ranged from 1 to 11. ΔK values were computed using the method of Evanno et al. (2005) in Structure Harvester. **B.** Log posterior likelihood of the runs plotted against the number of K clusters. All values were calculated from STRUCTURE runs with K ranging from 1 to 12, based on 20 replicate runs per K, with a burn-in period of 50,000 and 500,000 MCMC iterations.

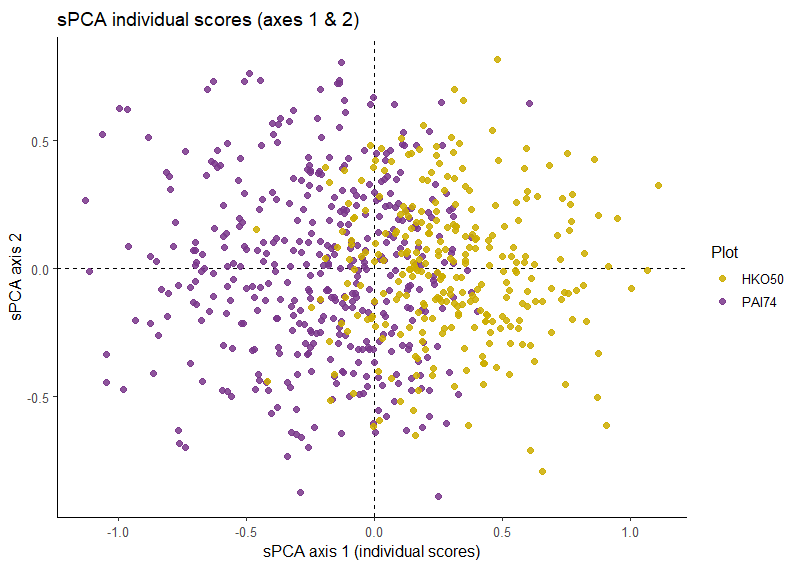

**Figure S4. Scatterplot of individual scores on the first two axes of the spatial Principal Component Analysis (sPCA).** Individuals from the two forest plots are represented by points coloured according to their plot of origin. Individuals from both plots overlap in the center of the graph, indicating a shared genetic background.

**Figure S5. Spatial patterns of genetic structure revealed by sPCA lagged scores in three study plots.** Interpolated maps of the first lagged principal component scores from spatial Principal Component Analysis (sPCA) for each study plot: (A) PAI74 and HKO50, (B) PAI74, and (C) HKO50. Black points indicate the locations of sampled individuals, and topographic isolines (5 m intervals) are overlaid to provide elevation context.
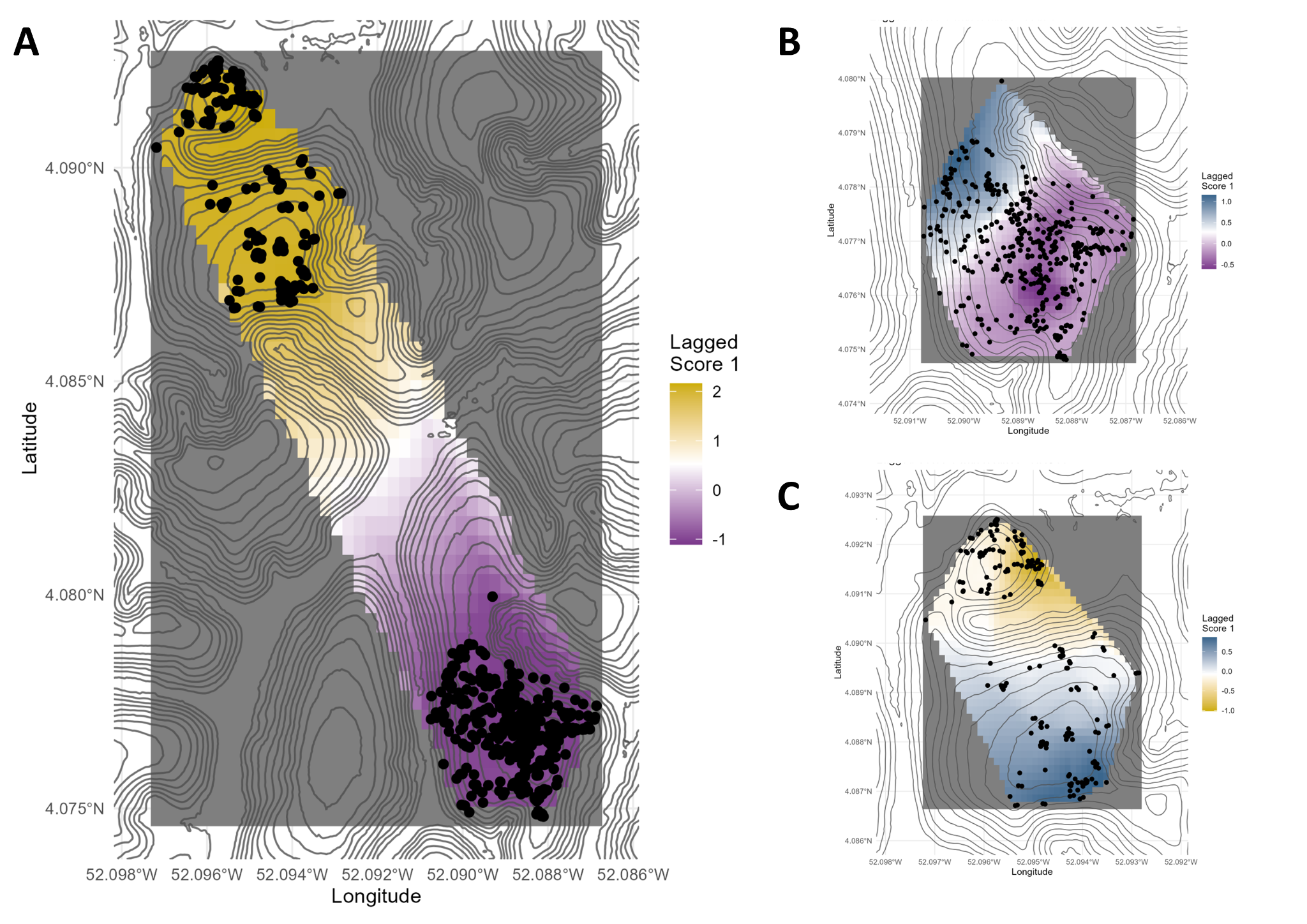

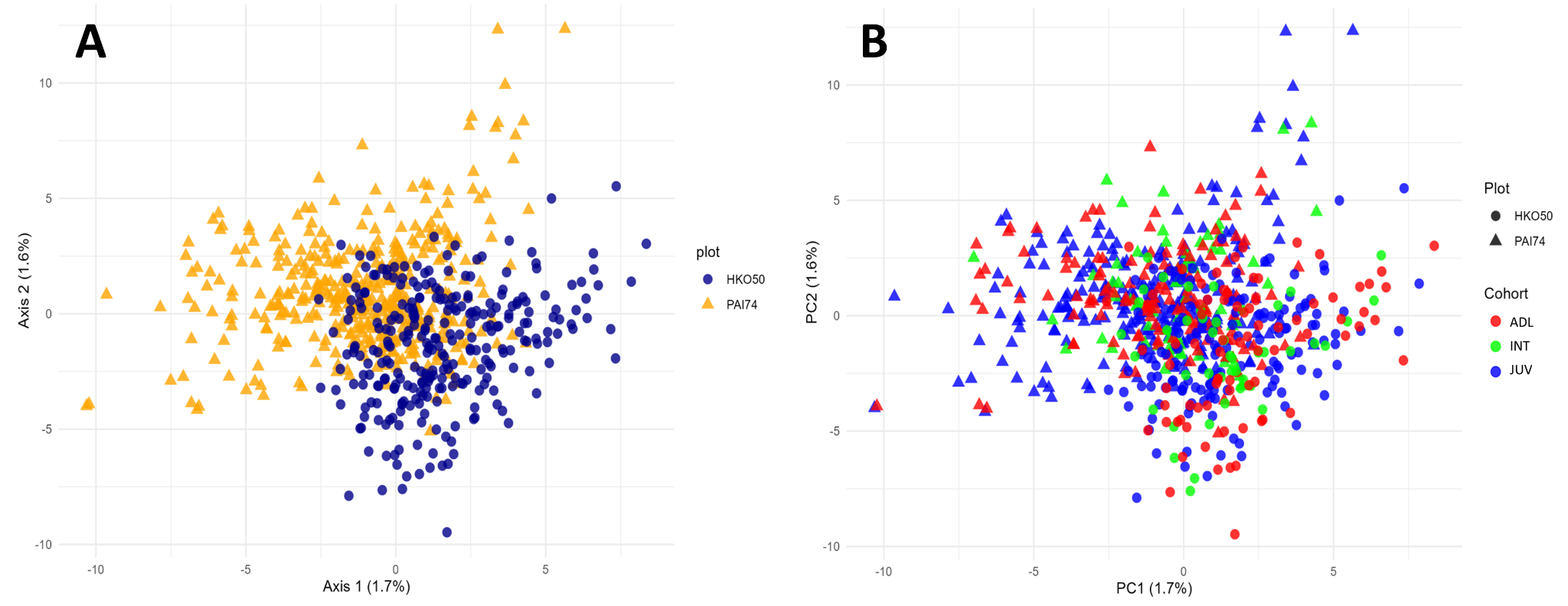

**Figure S6. Genetic differentiation and cohort structure across plots revealed by PCA. Principal Component Analysis (PCA) of nuclear microsatellite genotypes for D. guianensis individuals from the two study plots.** **A.** Individuals colored by plot, showing slight but distinguishable genetic differentiation between the plots. **B.** Same PCA with individuals colored by cohort: SED, INT, and ADL, with symbol shape indicating the plot of origin. The first two axes explain a combined 3.3% of total genetic variance.

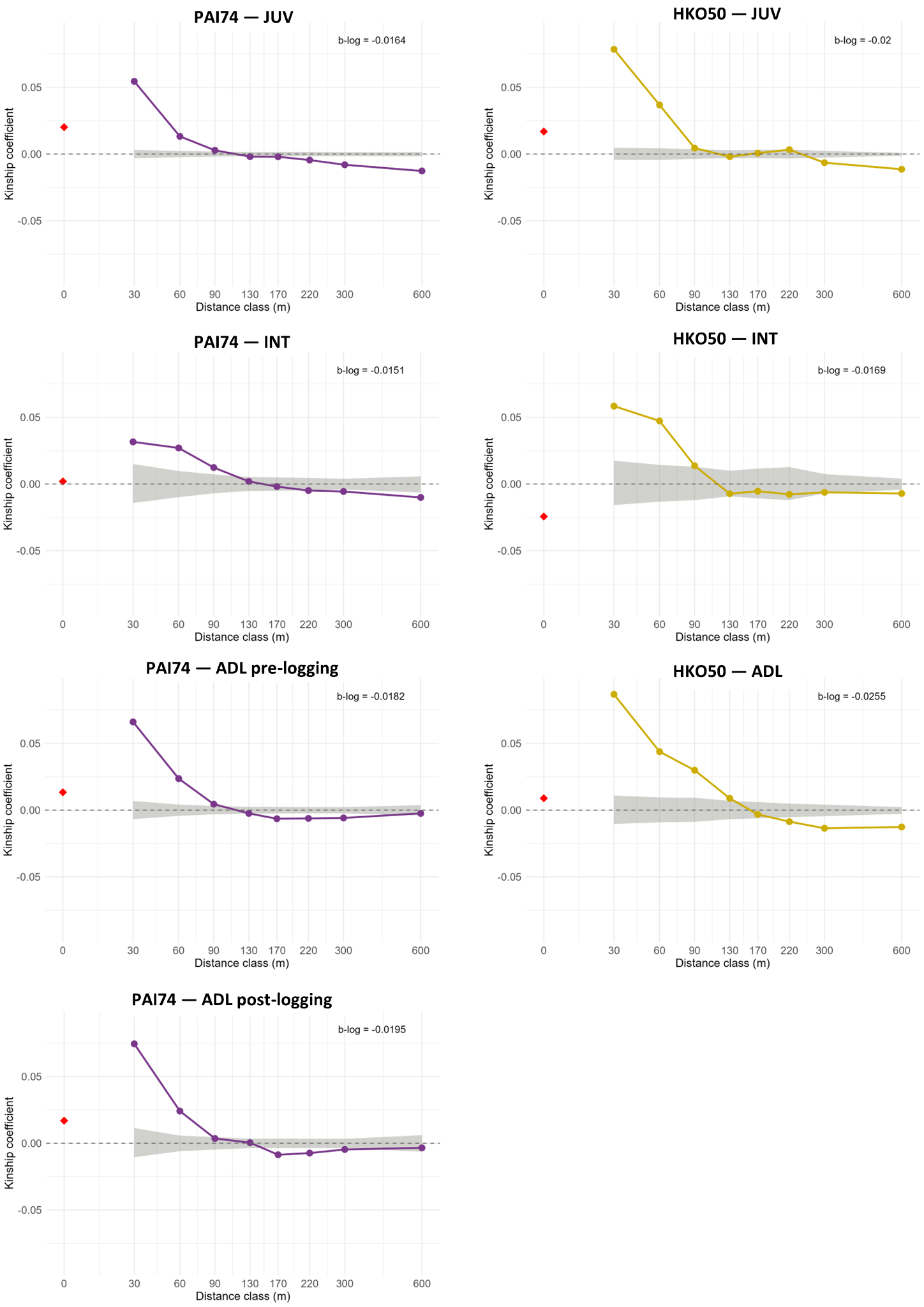

**Figure S7. Spatial autocorrelograms of the kinship coefficient (Fij) as a function of geographic distance for the two D. guianensis study plots for each of the three different cohorts (JUV, INT, ADL, pre-logging and post-logging).** The dashed grey area represents the 95% confidence interval for spatial randomness of genotypes, generated by 10,000 permutations of individual locations. The regression slope of kinship on log(distance) (b-log) is indicated for each plot. The red rhombus represents the intra-individual kinship coefficient.

**Figure S8. Distribution of seed and pollen dispersal distances in the logged plot PAI74 and the unlogged plot HKO50.** Histograms illustrate the frequency of effective dispersal events inferred from consistent parentage assignments (COLONY and CERVUS) in both study plots. Only the results of dispersions within plots are shown.
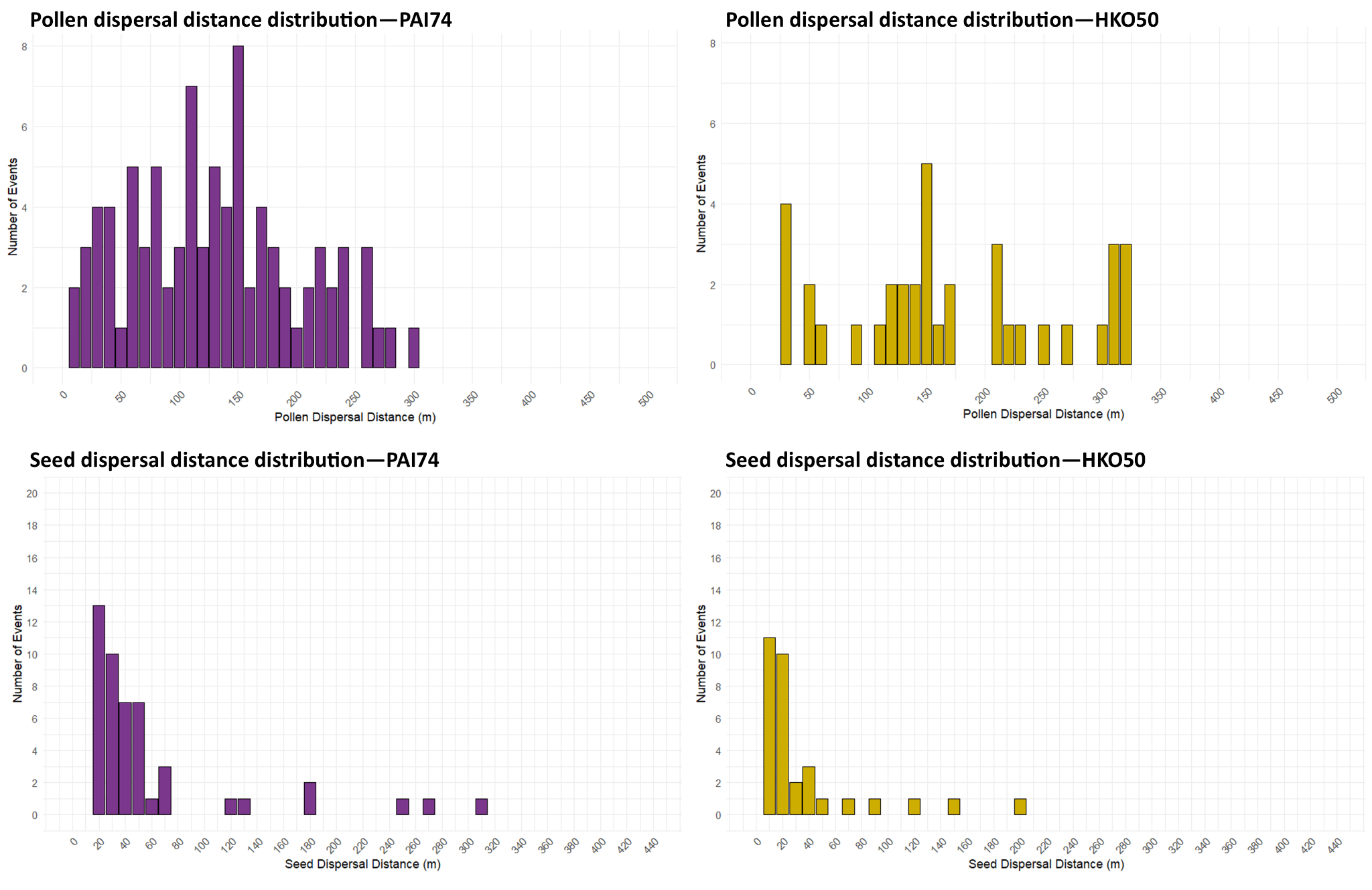

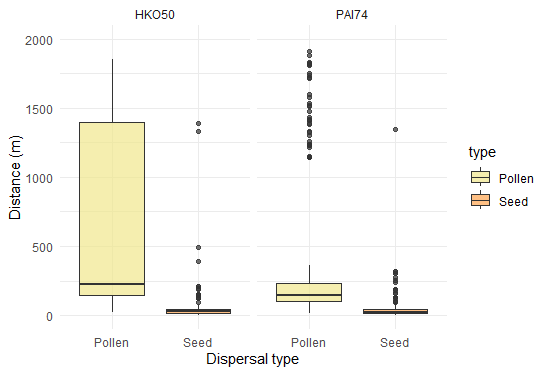

**Figure S9. Seed and pollen dispersal distances for offspring sampled in HKO50 and PAI74 plots with both parents identified.** Boxplots showing the distribution of dispersal distances mediated by seed (brown) and pollen (yellow) for offspring assigned to parents within each plot. In both plots, seed dispersal events are generally short-ranged, while pollen dispersal includes inter-plot dispersal and spans much larger distances, particularly in the unlogged plot HKO50. This contrast highlights the dominant role of pollen in long-distance gene flow, whereas seed dispersal contributes mainly to local recruitment. Note that this representation ignores dispersal from outside the plot, 143 seedlings in HKO50 and 157 in PAI74 had no both parents identified.

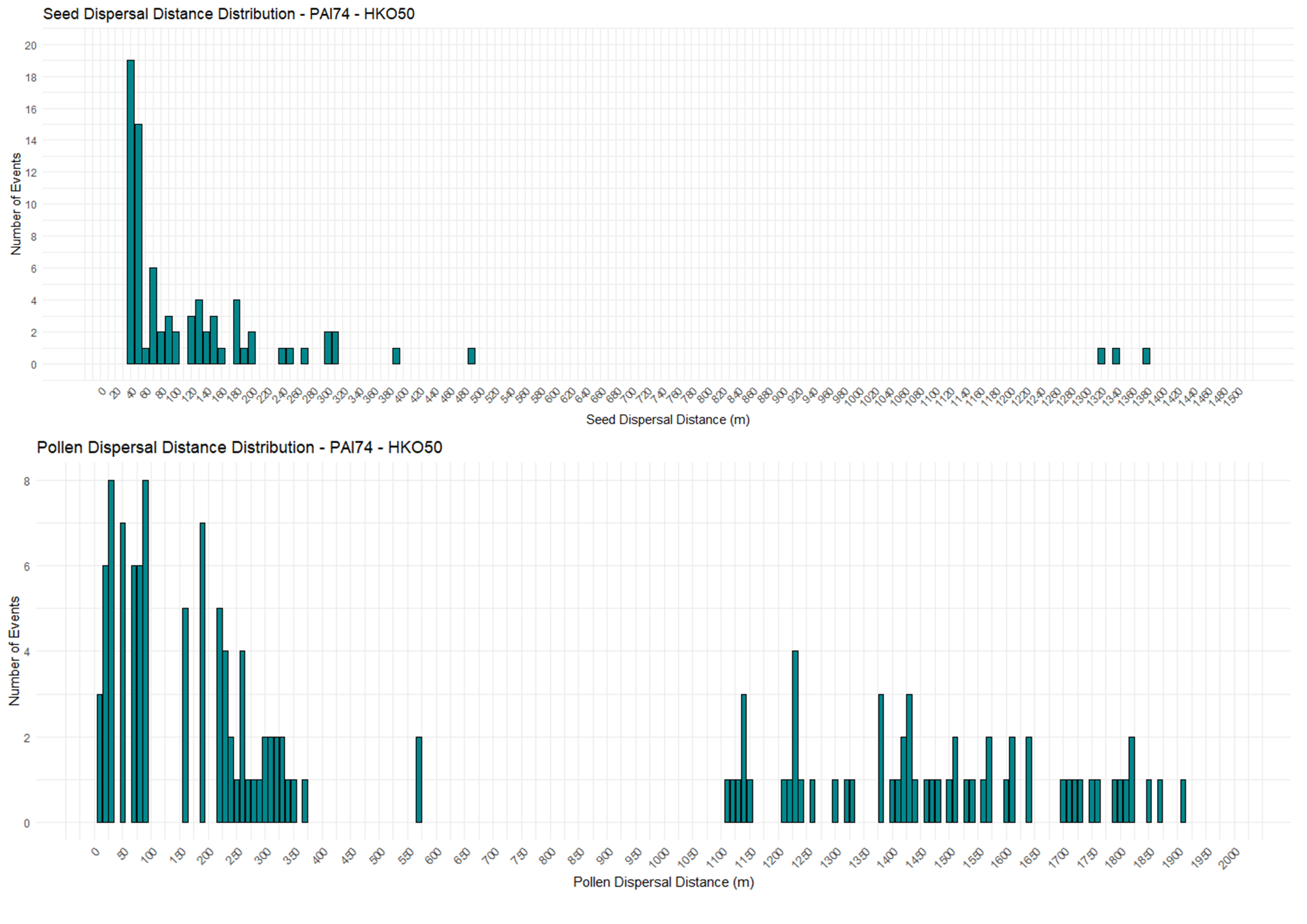

**Figure S10 Distributions of effective seed and pollen dispersal distances combining PAI74 (logged) and HKO50 (unlogged) plots.** Histograms illustrate the frequency of effective dispersal events inferred from consistent parentage assignments (COLONY and CERVUS).

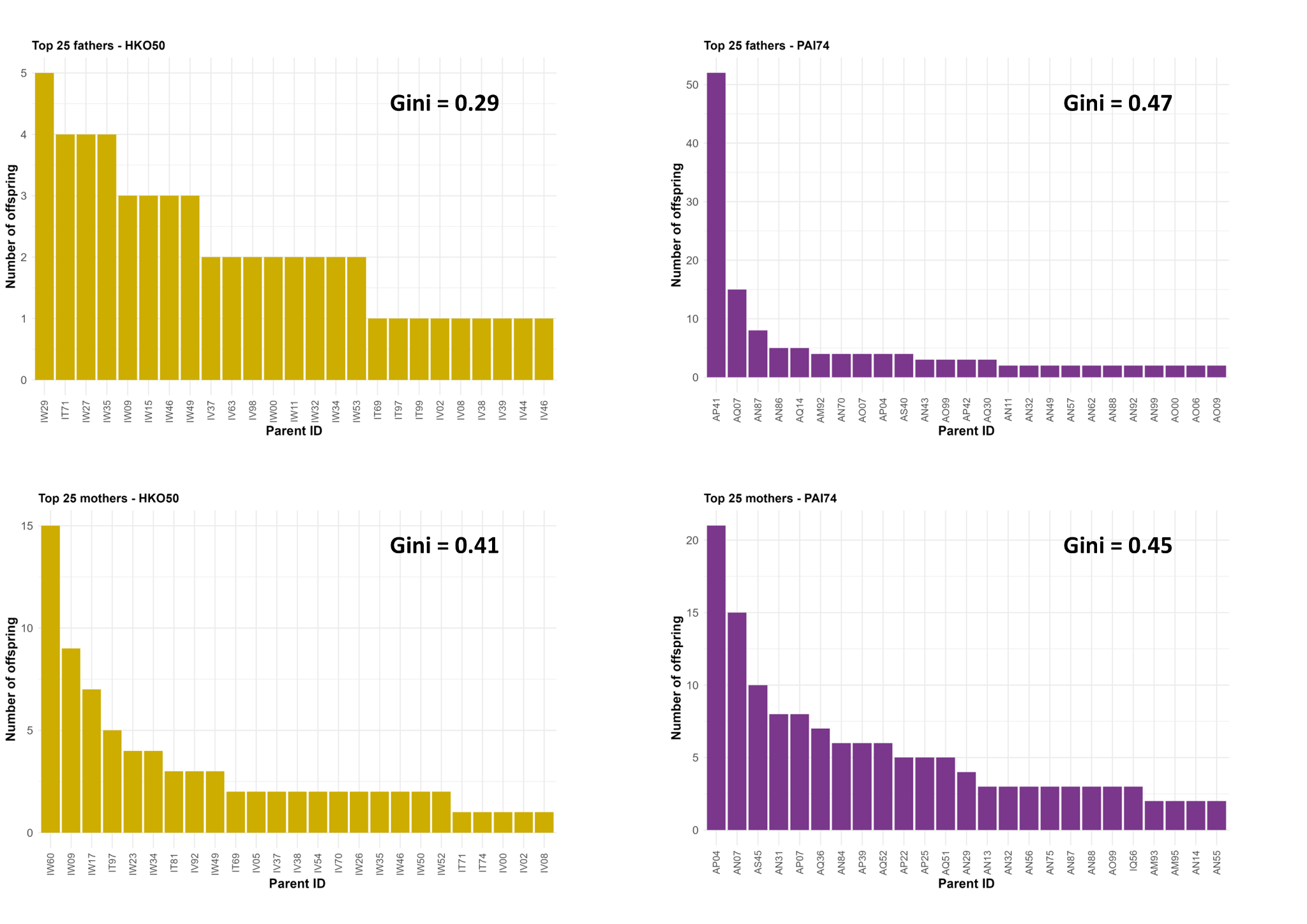

**Figure S11. Parental reproductive success and inequality in HKO50 and PAI74 plots.** Barplots showing the number of offspring assigned to the top 25 fathers and mothers in each plot. Bars are ordered by decreasing reproductive success, and parental IDs are shown on the x-axis. The Gini index, displayed in each panel, quantifies the degree of reproductive skew (0 = equal contribution, 1 = maximum inequality).
